# Abiotic stress-induced chloroplast and cytosolic Ca^2+^dynamics in the green alga *Chlamydomonas reinhardtii*

**DOI:** 10.1101/2024.06.21.600054

**Authors:** Matteo Pivato, Alex Costa, Glen Wheeler, Matteo Ballottari

**Author notes:** Corresponding author: Matteo Ballottari, Dipartimento di Biotecnologie, Università di Verona, Strada le Grazie 15, 37134 Verona Italy.

## Abstract

Calcium (Ca^2+^)-dependent signalling plays a well-characterized role in the perception and response mechanisms to environmental stimuli in plant cells. In the context of a constantly changing environment, it is fundamental to understand how crop yield and microalgal biomass productivity are affected by external factors. Ca^2+^ signalling is known to be important in different physiological processes in microalgae but many of these signal transduction pathways still need to be characterized. Here, the role of compartment-specific Ca^2+^ signalling was investigated in *Chlamydomonas reinhardtii* in response to environmental stressors such as nutrient availability, osmotic stress, temperature fluctuations and carbon sensing. An *in vivo* single-cell imaging approach was adopted to directly visualize signalling processes at the level of specific subcellular compartments, using *Chlamydomonas reinhardtii* lines expressing a genetically encoded ratiometric Ca^2+^ indicator. Hyper-osmotic shock caused cytosolic and chloroplast Ca^2+^ elevations, whereas high temperature and inorganic carbon availability primarily induced Ca^2+^ transients in the chloroplast. In contrast, hypo-osmotic stress only induced Ca^2+^ elevations in the cytosol. The results herein reported show that compartment-specific signalling pathways are likely to play an important role in the response of *Chlamydomonas* to these stimuli providing new understanding of the mechanisms exploited by microalgae to respond to specific natural conditions.

## Introduction

*Chlamydomonas* is worldwide distributed genus of unicellular microalgae that occupies a large diversity of ecological niches, including freshwater, marine and soil samples from temperate, tropical, and polar regions (E. H. Harris, 2013). *Chlamydomonas reinhardtii*, widely used as model organisms for unicellular green algae, have only been found in temperate soils in Northern America and Japan (Nakada, Shinkawa, Ito, & Tomita, 2010; ProLJschold, Harris, & Coleman, 2005; Sasso, Stibor, Mittag, & Grossman, 2018). Specifically, *C. reinhardtii* occurs mostly in cultivated fields, suggesting a preference for organic compounds and nutrient-rich soils. The ability to efficiently assimilate macronutrients, versatility in the tolerance of large variations in hydration or temperature, and the capability of using different sources of inorganic carbon, could all represent adaptive advantages in such environmental conditions.

Calcium (Ca^2+^)-dependent signalling mechanisms has a role in stress responses to environmental abiotic stimuli of *C. reinhardtii*, but also in many of its motile responses and in the regulation of photosynthesis (Bickerton, Sello, Brownlee, Pittman, & Wheeler, 2016; Verret, Wheeler, Taylor, Farnham, & Brownlee, 2010; Glen L. Wheeler & Brownlee, 2008). In a previous work *in vivo* single-cell imaging techniques were applied to study *C. reinhardtii* light-dependent Ca^2+^ signalling at subcellular resolution, reporting high light and/or H_2_O_2_ dependent chloroplast [Ca^2+^] transients (M. Pivato, Grenzi, Costa, & Ballottari, 2023). No cytosolic or mitochondrial Ca^2+^ responses to light stimuli were detected, leading to the hypothesis of a chloroplast-autonomous light-dependent Ca^2+^ response. However, chloroplast Ca^2+^ dynamics and their interaction with cytosolic Ca^2+^ transients still remain poorly understood in *C. reinhardtii*, together with the role of each compartment in the perception mechanisms of abiotic stress. Similarly, whether other environmentally relevant stimuli can result either in chloroplast and/or cytosolic compartment-specific [Ca^2+^] elevations has not been investigated so far. By studying how algae sense and adjust rapidly to the environment, it is possible to understand how they can cope with constantly changing natural conditions and gain new insights into the evolution of eukaryotic fundamental cellular processes.

In photosynthetic eukaryotes nitrogen (N) and phosphorous (P) sensing may require Ca^2+^ as a second messenger, even though distinct roles have been identified in different lineages. Nitrate resupply induces [Ca^2+^]_cyt_ elevations in N-limited *Arabidopsis* plants (K.-H. Liu et al., 2017; Riveras et al., 2015). Conversely, in diatoms P-limited cells exhibited previously unreported [Ca^2+^]_cyt_ elevations following phosphate resupply (Helliwell et al., 2021), which has not been observed in land plants. Whether N or P resupply leads to cytosolic or chloroplast [Ca^2+^] elevations in a green alga, or if the plant and diatom responses are conserved also in this organism is still not known.

Colonization of terrestrial environment during land plant evolution required adaptation to desiccation and high salinity of the soil. Since *C. reinhardtii* commonly inhabit temperate soil habitats, it needs to respond and adapt to substantial dehydration and salt challenges (Sasso et al., 2018). Hypo-osmotic stress, but not hyper-osmotic shifts, have been reported to evoke intracellular Ca^2+^ elevations in *C. reinhardtii* cells (Bickerton et al., 2016). However, osmotic Ca^2+^ signalling has not been investigated so far at the level of microalgal specific subcellular compartments, nor the relationship between cytosolic and chloroplast [Ca^2+^] has been explored in these responses yet. Plastidial-localized Ca^2+^ or K^+^ transporters in *Arabidopsis*, for instance, have a crucial role both in early steps and downstream signalling events that mediate osmotic stress responses, suggesting an important role of chloroplasts in this process (Stephan, Kunz, Yang, & Schroeder, 2016; Teardo et al., 2019).

Temperature fluctuations are also common environmental stressors encountered by plants and microalgae, and one of the main factors affecting their growth and productivity. During the last decades, the molecular mechanisms that regulate the responses to temperature stress in *C. reinhardtii* were deeply investigated (Ermilova, 2020; Schroda, Hemme, & Mühlhaus, 2015). High temperatures were also reported to rapidly induce ROS production in *C. reinhardtii* cells, H_2_O_2_ in cytosol and nucleus, and singlet oxygen in the chloroplast, potentially representing primary signalling molecules in the stress response (Niemeyer, Scheuring, Oestreicher, Morgan, & Schroda, 2021; Prasad, Ferretti, Sedlářová, & Pospíšil, 2016). However, the key sensor and/or the signal transducer molecules involved in the first steps of temperature sensing process are still uncharacterized. Likewise, the intracellular Ca^2+^ transient triggered by temperature stress has never been monitored in algae yet, to our knowledge. In plants, cold stress trigger transient cytosolic [Ca^2+^] elevations upon application, and interestingly heat stress stimulates chloroplast-specific [Ca^2+^] elevations (F. Gao et al., 2012; H. Knight, Trewavas, & Knight, 1996; Lenzoni & Knight, 2019). Recently, cytosolic [Ca^2+^] elevation in response to cold shock, but not to elevated temperatures, have also been reported in diatoms and suggested to have a role in short-term regulation of cellular processes, rather than longer term acclimation to a change in temperature (Kleiner et al., 2022).

In a previous work in *C. reinhardtii,* Ca^2+^ was also suggested to play a key role in the CO_2_ sensing mechanism (L. Wang et al., 2016). Since CO_2_ levels in aquatic environments can highly fluctuate and affect photoautotrophic growth (Maberly & Gontero, 2017), microalgae have evolved carbon concentrating mechanisms (CCMs) to elevate CO_2_ levels around Rubisco, the principal carbon-fixing enzyme. *C. reinhardtii* evolved a biophysical CCM that principally operates by the concentration of bicarbonate (HCO_3_^−^) inside the cell and its subsequent conversion to CO_2_ in the proximity of Rubisco, tightly localized in a microcompartment within the chloroplast, called pyrenoid (Y. Wang & Spalding, 2014). In addition, *C. reinhardtii* is also able to use the two-carbon molecule acetate either in the dark to support heterotrophic growth, or in the light, to support photoheterotrophic or mixotrophic growth (Elizabeth H Harris, 2001). Acetate is incorporated into acetyl-CoA and then mainly assimilated through the glyoxylate cycle (Lauersen et al., 2016; Wolfe Alan, 2005). Despite the biological relevance of carbon fixation in nature, how photosynthetic cells sense and respond to carbon availability remains poorly understood.

Ca^2+^ signalling is implicated in regulating the transition between Ci uptake pathways at ambient or sub-ambient CO_2_ in *C. reinhardtii*: under CO_2_-limiting conditions [Ca^2+^] is elevated in the pyrenoid and the Ca^2+^-binding protein (CAS) relocalizes from the stroma to the pyrenoid tubules (L. Wang et al., 2016). Pyrenoid [Ca^2+^] controls the expression of a set of CCM genes by CO_2_-regulated Ca^2+^-dependent retrograde signalling to the nucleus (L. Wang et al., 2016). Elevated [Ca^2+^] within the pyrenoid was detected under low CO_2_ conditions using the fluorescent dye Calcium Green-1 (Wang et al., 2016). However, care must be taken when examining sub-cellular Ca^2+^ using a single-wavelength Ca^2+^-sensitive dyes that have been loaded as acetoxymethyl ester derivatives, as these can display unequal loading and dye compartmentalization in plant and algal cells (Braun & Hegemann, 1999; Hong-Hermesdorf et al., 2014). To overcome this limitation and distinguish between higher dye abundance in the pyrenoid and *bona fide* pyrenoid [Ca^2+^] elevations, chloroplast targeted ratiometric Ca^2+^ indicators are needed. Current knowledge about the role of Ca^2+^ in the functioning of the CCM in *C. reinhardtii* is somewhat fragmented and the physiological significance of its dynamics in limiting CO_2_ conditions is not yet clear. Moreover, real-time measurements of intracellular Ca^2+^ dynamics in the responses to inorganic carbon (Ci) availability have never been reported in this organism to our knowledge.

Here, intracellular Ca^2+^ transients at the level of different subcellular compartments in responses to nutrients availability, osmotic stresses, temperature variations and/or organic or inorganic carbon availability were investigated in *C. reinhardtii* by using YC3.6 ratiometric Ca^2+^ indicator, expressed in the cytosol or in the chloroplast. The objective of this investigation is to assess the role of Ca^2+^ in sensing different environmental stimuli and the contribution of different compartments (cytosol and chloroplast) to these signalling pathways, determining whether there is conservation or differentiation of the Ca^2+^ signalling responses between green algae and land plants.

## Materials and Methods

### Algal strains and culture conditions

The *C. reinhardtii* strains used in this study were UVM4 (UV-mediated mutant 4) (Neupert, Karcher, & Bock, 2009) and the previously obtained transgenic UVM4 *C. reinhardtii* lines expressing YC3.6 (Nagai, Yamada, Tominaga, Ichikawa, & Miyawaki, 2004) at the level of the cytosol or the chloroplast stroma (M. Pivato et al., 2023). Algal cells were cultivated in mixotrophic or photoautotrophic conditions, in Tris-acetate-phosphate (TAP) or Tris-phosphate (TP) minimal medium respectively (E. H. Harris, 2008; Kropat et al., 2011). Liquid cultures were maintained in shake flasks, whereas agar-solidified medium in plates, at 18°C and 70 μmol photons m^−2^ s^−1^ on 16:8 h (light:dark) circadian cycle, unless otherwise stated. Cultures were maintained in the exponential growth phase. For the Ca^2+^ free (-Ca^2+^) TAP or TP media used, the same recipe for TAP or TP was used, but without CaCl_2_ and adding 200 μM EGTA.

For the nutrient (N, P) limitation treatments for nutrient resupply experiments, cells were grown in TAP or TP medium, but with concentrations of ammonium (NH_4_Cl) or phosphate (K_2_HPO_3_ and KH_2_PO_4_) reduced to one tenth or one hundredth respectively of those typically found in standard medium recipes (0.75 mM and 10 μM of ammonium and phosphate, respectively).

### Confocal Laser Scanning Microscopy

Confocal Laser Scanning Microscopy (CLSM) analyses were performed using a Leica SP8 or SP5 confocal microscope and LASX (Leica, Wetzlar, Germany) application suite for acquisition and analysis. Images were acquired by a 63X 1.40 NA oil immersion objective with different digital zoom. Chloroplast were visualized using chlorophyll autofluorescence.

Ca^2+^ imaging of living cells was performed as described previously (Loro & Costa, 2013). YC3.6 was excited at 458 nm, and emission of FRET pair proteins ECFP and cpVenus was collected at 475-505 nm and 525-545 nm, respectively, with 2 Hybrid spectral detectors. Chlorophyll autofluorescence was excited with 458 nm and detected at 680-720 nm.

Images were analyzed using ImageJ software. cpVenus and CFP emissions of the analyzed regions of interest were used for the FRET ratio calculation (cpVenus/CFP) and, where suitable, normalized to the initial ratio and plotted versus time. Background subtraction was performed independently for both channels before calculating the ratio. Maximal FRET Ratio variation was calculated subtracting the median of the pre-stimulus FRET Ratio values.

### Chlamydomonas reinhardtii cells imaging

Mid/Late-log phase *C. reinhardtii* cell cultures were kept in light until the experiment, when they were placed into a 35 mm glass-bottomed dish (*In Vitro* Scientific, Sunnyvale, CA, USA) coated with 0.05% poly-L-lysine (Sigma-Aldrich, St Louis, MO, USA) to facilitate adherence of the cells. Cells were perfused with their culturing medium with a gravity driven perfusion system at a flow rate of 5 mL min^-1^. Each specific stimulus was delivered by rapidly switching the perfusion from the culturing medium to the culturing medium + stimulus. Hyper-osmotic shocks were delivered by switching the perfusion to TAP + 100-300 mM NaCl or 360 mM Sorbitol. Hypo-osmotic treatments were delivered by switching the perfusion to deionized water containing 340 μM CaCl_2_. To investigate the response to hypo-osmotic shock in the absence of contractile vacuole activity, cells were equilibrated to TAP containing 100 mM sorbitol for at least 1h. The N and P resupply treatments were delivered by switching the perfusion from TAP or TP medium without ammonium or phosphate respectively, to the standard TAP or TP medium. Heat and cold shocks were delivered by switching the perfusion to the same medium at the desired different temperatures, measured directly in the perfusion chamber during experiments through a digital infrared thermometer (10°C for cold and 38, 41 and 46°C for heat stress). The same set-up was used for carbon sensing experiments, except the perfusion was switched from TP to TP + 10-50 mM NaHCO_3_ or TAP (TP + 17 mM acetate). Steady-state FRET Ratio was monitored illuminating the cells by continuous laser light at 458 nm for 2 minutes before each measurement, to ensure signal stability.

### Protein extraction and SDS-PAGE

Protein extracts were analyzed by SDS-PAGE as described in Laemmli (Laemmli, 1970). Western blot analysis was performed using anti-CAH3 and anti-RbcL antibodies (Agrisera Antibodies) and an anti-rabbit Immunoglobulin G Alkaline Phosphatase-conjugated secondary antibody (Merck).

### Statistical Analysis

Students’ two-tailed *t* test or One-way ANOVA for independent samples were applied to statistically evaluate results. Error bars indicate standard deviation (SD). Significance thresholds and the number of replicates used to calculate SD are indicated in the figure legends.

## Results

### Intracellular [Ca^2+^] elevations are not observed following nutrient resupply in C. reinhardtii cells

To investigate the role of Ca^2+^ signalling in nutrient sensing in *C. reinhardtii*, the previously obtained transgenic *C. reinhardtii* lines, encoding the ratiometric genetically encoded fluorescent Ca^2+^ indicator, YC3.6, at the level of the cytosol and chloroplast stroma (M. Pivato et al., 2023) were exploited. Similar experimental approach to investigate Ca^2+^-dependent nutrient sensing mechanisms previously reported in diatoms (Helliwell et al., 2021) was here applied in *C. reinhardtii.* YC3.6 expressing cells were grown in mixotrophic or photoautotrophic conditions, respectively in TAP or TP medium, but with reduced concentrations of ammonium or phosphate. Optimal nutrient concentrations inducing starvation and growth impairment was preliminarily determined by growing cells in different ammonium or phosphate regimes in mixotrophic (Figure 1a, d) and photoautotrophic (a, d) conditions: (1) replete condition (7.5 mM and 1 mM respectively); (2) limited condition (0.75 mM and 10 µM respectively). Both limited condition treatments significantly impaired cell growth compared to replete conditions (Figure 1a, d). According to the results obtained, cells for the following analysis were sampled after 52 and 76 hours for the mixotrophic growth in ammonium limited conditions (Figure 1a), and after 76 and 100 hours for the mixotrophic growth in phosphate limited conditions (Figure 1d). Single cell *in vivo* [Ca^2+^] at the level of the cytosol and chloroplast were monitored in nutrient deplete YC3.6 expressing cells, following resupply with each respective nutrient. No relevant changes of the cytosolic or chloroplast [Ca^2+^] were detected following perfusion with TAP medium containing ammonium or phosphate restored to 7.5 mM (Figure 1b, c) and 1 mM (Figure 1e, f) respectively. No response was detected also in photoautotrophic grown nutrient depleted cells, where the same experimental approach was applied. From these experimental analysis no evidence for a role for Ca^2+^ signalling in sensing exogenous ammonium or phosphate could be detected, differently from what has been observed in diatoms or plants (Helliwell et al., 2021; K.-H. Liu et al., 2017; Riveras et al., 2015). The reported experiments may thus exclude the presence of a conserved cytosolic or chloroplast Ca^2+^-dependent signalling pathway for environmental phosphate or ammonium sensing in *C. reinhardtii* in the tested conditions. However, we cannot rule out that the lack of any observable change in the Ca^2+^ concentrations might depend on the lower sensitivity of YC3.6 compared to the other Ca^2+^ indicators used in previous works (R-GECO1 in Helliwell et al., 2021; aequorin in Riveras et al., 2015; GCaMP6s in Liu et al., 2017).

**Figure 1.**
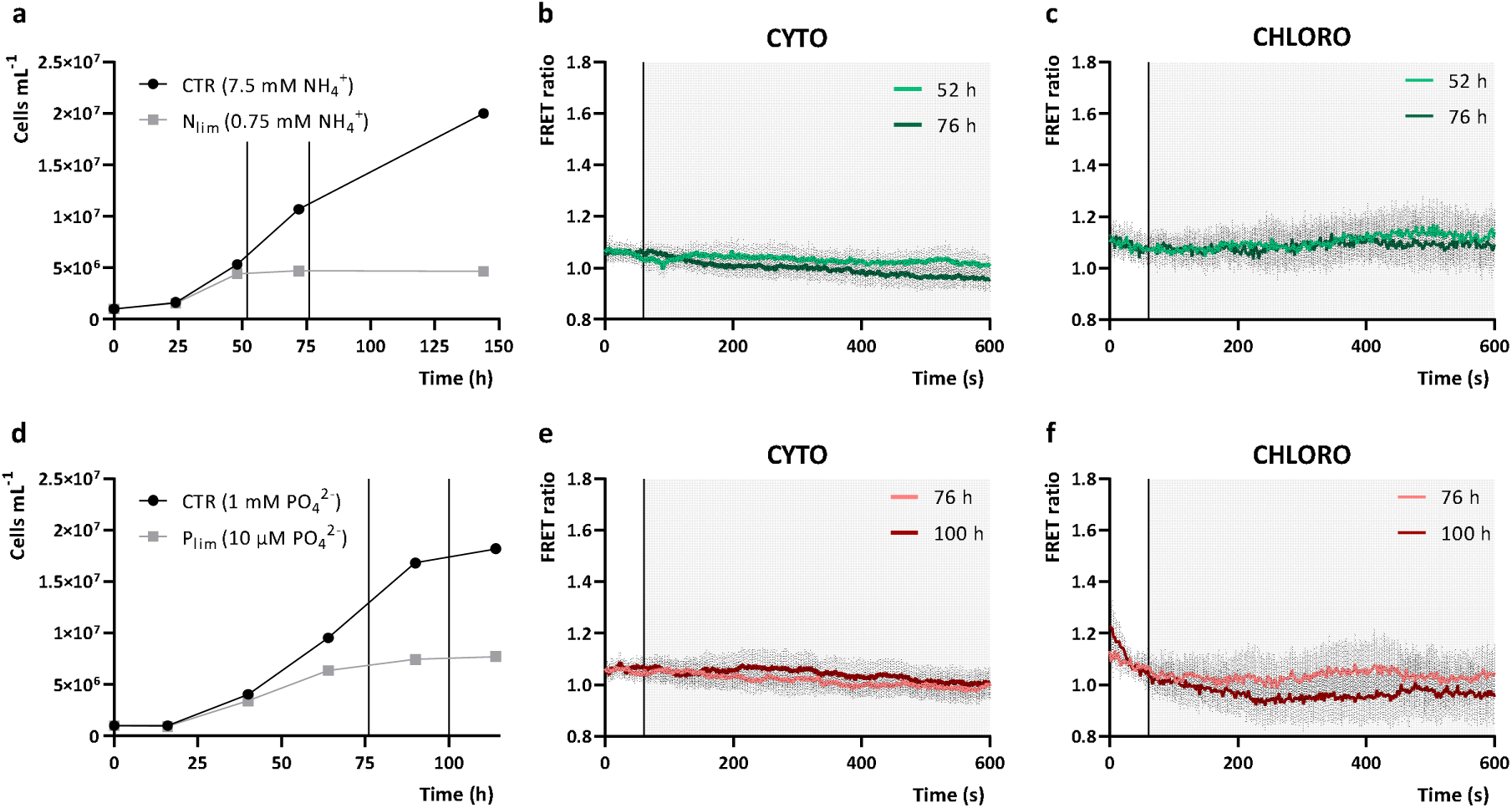
YC3.6 FRET ratio of nutrient-limited *C. reinhardtii* cells in response to ammonium or phosphates resupply. a, d. Mixotrophic growth over time of *C. reinhardtii* cells in standard TAP medium with ammonium-(a) or phosphate-replete (d) (CTR, 7.5 mM NH_4_^+^ and 1 mM PO_4_^2-^ respectively), versus ammonium- or phosphate-limited (N_lim_ 0.75 mM NH_4_^+^ and P_lim_ 10 µM PO_4_^2-^ respectively) conditions (n = 1). Black vertical lines in the graphs indicate the time-points of the growth curve at which nutrient deplete YC3.6 expressing cells were harvested to perform nutrient resupply experiments (reported in different colour shades in b, c, e and f). b, c. Averaged and normalized FRET Ratio ± SD of cytosolic (b, n > 36 cells) and chloroplast (c, n = 15) YC3.6 in *C. reinhardtii* cells in response to ammonium resupply (TAP, 7.5 mM NH_4_^+^) (black vertical line indicate the onset of the stimulus, at 60s). e, f. Averaged and normalized FRET Ratio + SD of cytosolic (e, n > 39 cells) and chloroplast (f, n > 13) YC3.6 in *C. reinhardtii* cells in response to phosphate resupply (TAP, 1 mM PO_4_^2-^) (black vertical line indicate the onset of the stimulus, at 60 s).

### Hyper-osmotic stress induces cytosolic and chloroplast [Ca^2+^] elevations

To investigate whether Ca^2+^ signalling processes are conserved between vascular plants and green algae, *C*. *reinhardtii* YC3.6 expressing lines were subjected to osmotic stress known to induce [Ca^2+^] elevations in the cytosol and chloroplast compartments of land plants cells (Sello et al., 2018; Sello et al., 2016). Salt stress has already been reported to evoke cytosolic Ca^2+^ elevations in *C. reinhardtii* cells (Bickerton et al., 2016). Accordingly, *C. reinhardtii* cells treatment with increasing sodium chloride (NaCl) concentrations (100-300 mM NaCl) induced significant cytosolic [Ca^2+^] transients. Importantly, the same stress also induced a chloroplast [Ca^2+^] increase, with different dynamic and kinetic parameters (Figure 2). Treatment of the cells with incremental concentrations of NaCl (100, 200 and 300 mM) resulted in progressive increases in the cytosolic maximal FRET Ratio variation (Figure 2c), indicating that the amplitude of NaCl-induced cytosolic [Ca^2+^] elevations are sensitive to the intensity of the stimulus. A significant shrinkage of the cells was also observed immediately after the end of the stimulus with all the NaCl treatments applied (Figure 2a, b); the magnitude of cell area decrease is related to the intensity of the applied stimulus and indicates a clear shrinkage effect of the salt stress to *C. reinhardtii* cells. NaCl-induced chloroplast [Ca^2+^] elevations were instead observed only with 200 mM and 300 mM NaCl treatments (Figure 2d), suggesting the presence of a threshold stimulus required to trigger chloroplast [Ca^2+^] elevations. Higher concentrations of NaCl (≥ 400 mM) shrank the cells excessively, preventing accurate determination of [Ca^2+^] in both the compartments. Interestingly, iso-osmotic sorbitol treatments induced comparable compartment-specific [Ca^2+^] increases in both the cytosolic and chloroplast compartment (Figure 2e, f), pointing out the osmotic effect as the major responsible for the monitored responses. Moreover, the depletion of extracellular [Ca^2+^] did not alter the hyper-osmotic-induced Ca^2+^ signature in both the subcellular compartments (Figure 2e, f), suggesting a relevant role of intracellular compartments as main Ca^2+^ sources.

**Figure 2.**
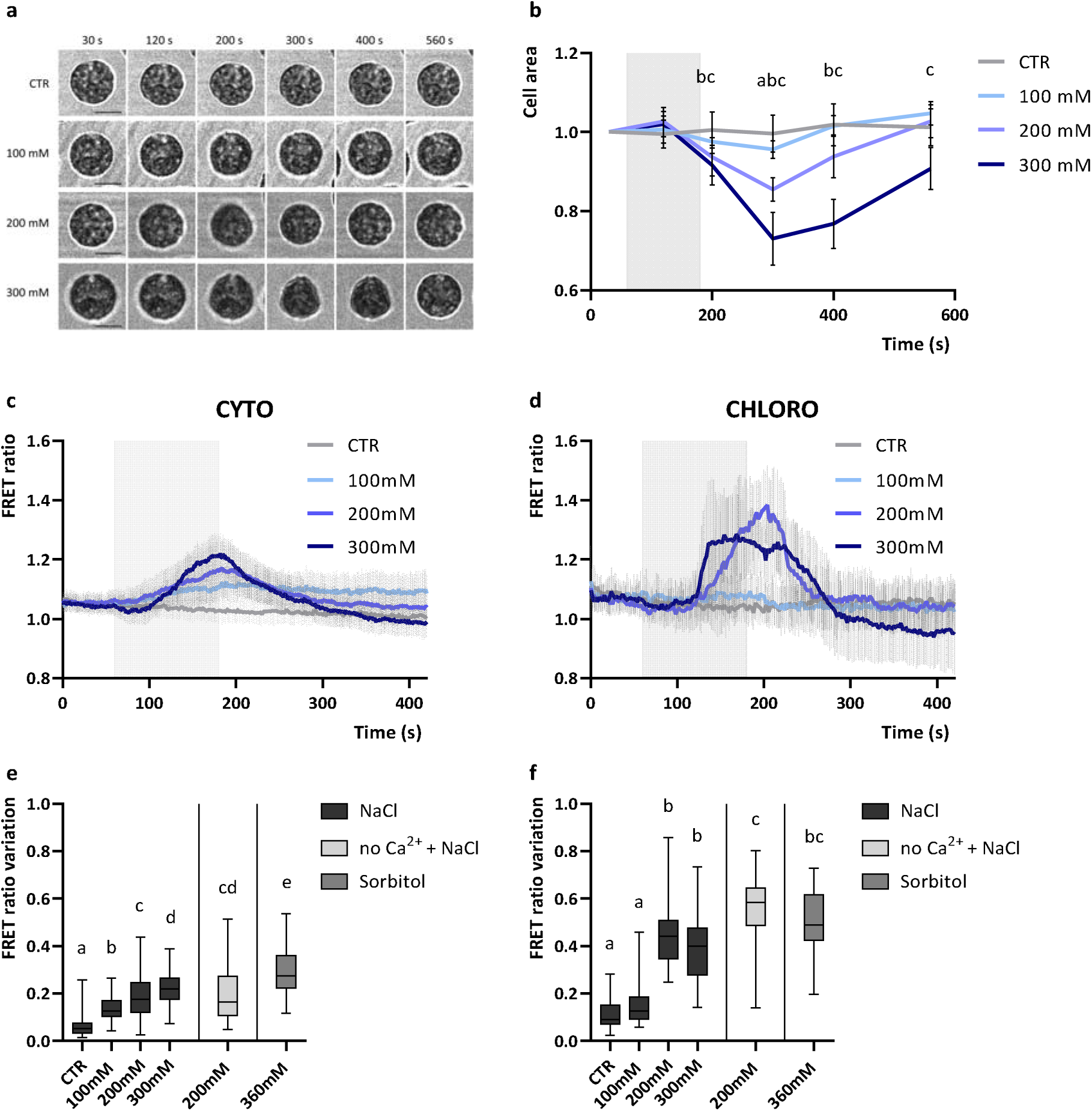
Cytosolic and chloroplast Ca^2+^ dynamics and cell area variation of *C. reinhardtii* cells in response to hyper-osmotic stimuli. a. Representative images of *C. reinhardtii* cells in response to hyper-osmotic stimuli at different time points (referred to NaCl treatment in b., 120 s of switch from growth medium to growth medium + X mM NaCl at 60 s, where X is 100, 200 or 300 mM). Scale bars: 5 µm. b. Averaged and normalized cell area measurements ± SD of cell area (n = 15 cells) in response to hyper-osmotic stimuli (grey rectangle indicate the treatment, 120 s of switch from growth medium to growth medium + X mM NaCl, where X is 100, 200 or 300 mM). Significantly different values from the control (untreated cells, CTR) are marked with different letters indicating the treatment (a, 100 mM NaCl; b, 200 mM NaCl; c, 300 mM NaCl) as determined by One-way ANOVA (P < 0.05). c, d. Averaged and normalized FRET Ratio ± SD of cytosolic (c, n > 35 cells) and chloroplast (d, n > 19) YC3.6 in C. *reinhardtii* cells in response to hyper-osmotic stimuli (grey rectangle indicate the treatment, 120 s of switch from growth medium to growth medium + X mM NaCl). e, f. Maximum FRET Ratio variations triggered by hyper-osmotic stimuli respectively in the cytosol (e, n > 35 cells) and chloroplast (f, n > 19 cells). One-way ANOVA: P value < 0,05.

The spatial characteristics of the hyper-osmotic-induced chloroplast [Ca^2+^] elevations were then examined. Upon 300 mM NaCl treatment, *C. reinhardtii* cells shrank, rapidly changing chloroplast shape (Figure 3a, d), while chloroplast [Ca^2+^] proportionally rose in the location of the pyrenoid (Figure 3b, c, e, g), a microcompartment situated in the center of the cup-shaped chloroplast and known to promote photosynthetic CO_2_ fixation by the enzyme ribulose-1,5-bisphosphate carboxylase/oxygenase (Rubisco) (Barrett, Girr, & Mackinder, 2021). Interestingly, the pyrenoid resting steady-state FRET ratio before the stimulus was significantly higher compared to the other chloroplast regions, indicating higher free Ca^2+^ concentrations (Fig. 3f). This is in accordance with what was previously observed in both high- and low-CO_2_ acclimated cells, suggesting higher free Ca^2+^ concentrations in the pyrenoid region (L. Wang et al., 2016). The cell volume alterations following the hyper-osmotic shocks, together with the transient increase of intracellular [Ca^2+^], might indicate the activation of mechanosensitive and/or osmosensitive ion channels, both at the level of the plasma membrane and the chloroplast envelope or thylakoid membranes.

**Figure 3.**
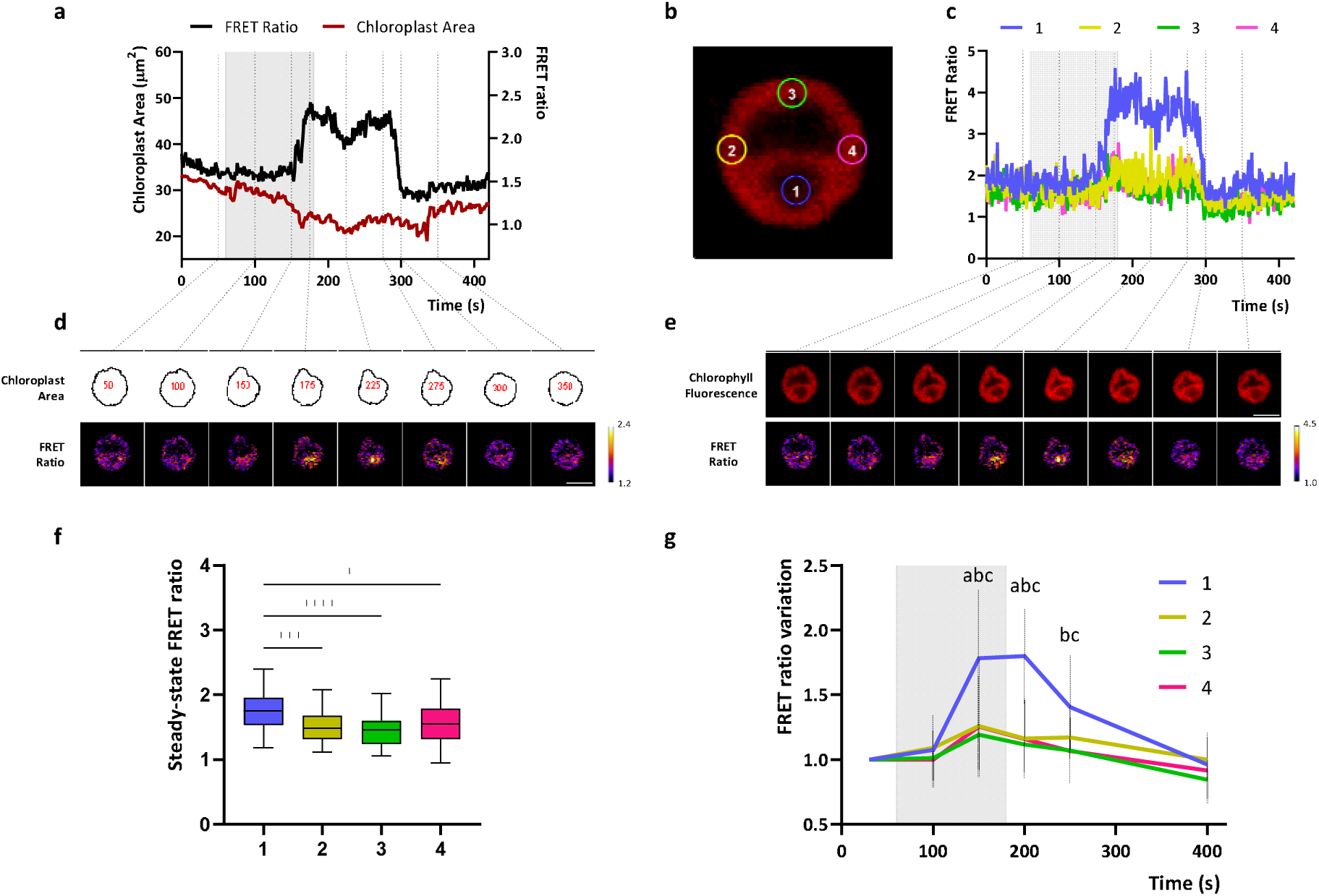
Spatial characterization of a chloroplast [Ca^2+^] elevation induced by hyper-osmotic shock in a representative *C. reinhardtii* cell. a. Normalized FRET Ratio trace and chloroplast area variation of a single representative cell upon hyperosmotic stimulus (200 mM NaCl, grey rectangle). Below (d.) are reported example images at the indicated time-points of the chloroplast area and pseudocoloured images of FRET Ratio signal. c. FRET Ratio trace and chlorophyll fluorescence of a single representative cell (same of a.) upon hyper-osmotic stimulus (300 mM NaCl, grey rectangle). In different colors are reported the FRET Ratio traces analyzed from specific regions of interest within the chloroplast (drawn and referred to b.). Below (e.) are reported example images at the indicated time-points of the chlorophyll fluorescence and pseudocoloured images of FRET Ratio signal. Scale bars (white line): 5 µm. f. Basal steady-state FRET ratios at the level of the different subchloroplast regions reported in b. (n□=□40 cells). One-way ANOVA: ****, P□<□0.0001; ***, P□<□0.001; *, P< 0.05. g. Averaged and normalized chloroplast FRET ratio variations ± SD (n = 15 cells examined where the pyrenoid region could be clearly distinguished) in response to hyper-osmotic stimuli (grey rectangle indicate the treatment, 120 s of switch from growth medium to growth medium + 300 mM NaCl). Significantly different values from the pyrenoid region (1, blue line) are marked with different letters indicating the region (a, 2; b, 3; c, 4) as determined by One-way ANOVA (P□<□0.05).

### Hypo-osmotic stress induces cytosolic [Ca^2+^] elevations

Hypo-osmotic shock applied by switching from TAP to deionized water (containing 0.34 mM CaCl_2_) did not affect cytosolic [Ca^2+^] (2a), as previously reported for cell wall-deficient strains (Bickerton et al., 2016). However, cells acclimated to a higher osmolarity (TAP + 100 mM sorbitol) displayed hypo-osmotic induced cytosolic [Ca^2+^] elevations when perfused with deionised water (+ 0.34 mM CaCl_2_), with a significantly higher maximal FRET Ratio variation (2b, c). These cells showed a significant increase in the cell area in response to the hypo-osmotic stimulus, whereas TAP acclimated cells showed no significant difference compared to control (Figure S3a). The results obtained show that 23.4% of the cells exhibited cytosolic [Ca^2+^] elevations (n = 14) in the form of brief spikes, defined as 0.2-point variation of the maximal FRET Ratio value. The spatial and temporal characteristics of these transients are clearly different to the hyper-osmotic induced sustained cytosolic [Ca^2+^] increases (Fig. 2c), but comparable to the previously reported hypo-osmotic induced cytosolic [Ca^2+^] elevation in the wild-type walled strain CC1021 (Bickerton et al., 2016). Both hypo-osmotic shocks (deionized water + 0.34 mM CaCl_2_ to cells acclimated either to TAP or to TAP + 100 mM sorbitol) did not trigger chloroplast-specific [Ca^2+^] transients during the hypo-osmotic stress (2d, e). Significant chloroplast [Ca^2+^] elevations occurred when the perfusion was restored to the initial medium (TAP or TAP + 100 mM sorbitol) (Figure S2f). As hyperosmotic shocks trigger chloroplast [Ca^2+^] elevations (Figure 2c, d), these findings suggest that the cells rapidly acclimate to the lower osmolarity of the hypo-osmotic stimulus, resulting in a hyper-osmotic shock when the perfusion is restored to the initial medium. In support of this, a significant decrease of the cell area was detected when the perfusion was restored to TAP medium after the hypo-osmotic shock (Figure S3a). However, cells acclimated to a higher osmolarity (TAP + 100 mM sorbitol) do not show an immediate decrease in area when they are restored to their original media. These cells undergo rapid expansion during the hypo-osmotic shock, which slows immediately when returned to TAP + 100 mM sorbitol. It is possible that there is a contraction of the chloroplast volume within the expanding cell wall in these cells that we are unable to detect by light microscopy. The results obtained suggest the signalling response to the hypo-osmotic shock is specific to the cytosol and represent a series of brief spikes rather than a sustained [Ca^2+^] elevation.

### Heat stress induces chloroplast-specific [Ca^2+^] elevations

In plants both cold and heat stress differentially trigger transient intracellular [Ca^2+^] elevation upon application, with heat stress stimulating chloroplast free [Ca^2+^] elevations and cold stress affecting cytosolic [Ca^2+^] (Lenzoni & Knight, 2019). Similarly, in diatoms cold shock has been reported to trigger cytosolic [Ca^2+^] elevations (Kleiner et al., 2022). To test the evolutionary conservation of these mechanisms in green algae, perfusion experiments with the culturing medium at high or low target temperatures were performed with *C. reinhardtii* cytosolic and chloroplast YC3.6 expressing lines. Actual temperatures in the perfusion chamber were recorded and are displayed in the figures for all experiments (Figure 4, S4); it should be noted that they differed by ± 2°C from the initial medium temperatures, due to equilibration of the small volume of warm or cold perfusate with room temperature (RT). It should be considered that fluorescence itself and excited-state lifetimes of calcium dyes are strongly affected by temperature, even if no specific information are available in the case of YC3.6 indicator (Oliver, Baker, Fugate, Tablin, & Crowe, 2000). If cpVenus and CFP fluorophores do not respond equally to temperature, large changes in temperature could transiently influence YC3.6 fluorescence properties during the stimulation phase, without affecting its subsequent functionality. We found that temperature treatments on YC3.6 expressing cells caused a slight change of the FRET ratio during the stimulus in all the tested conditions (positive during heat stress and negative in cold shocks), limiting our ability to resolve potential [Ca^2+^] elevations during temperature shocks themselves. We therefore examined whether the temperature stress resulted in sustained [Ca^2+^] elevations, either in the cytosol or chloroplast, by monitoring cells after the perfusion chamber temperature was restored to the initial value (RT).

**Figure 4.**
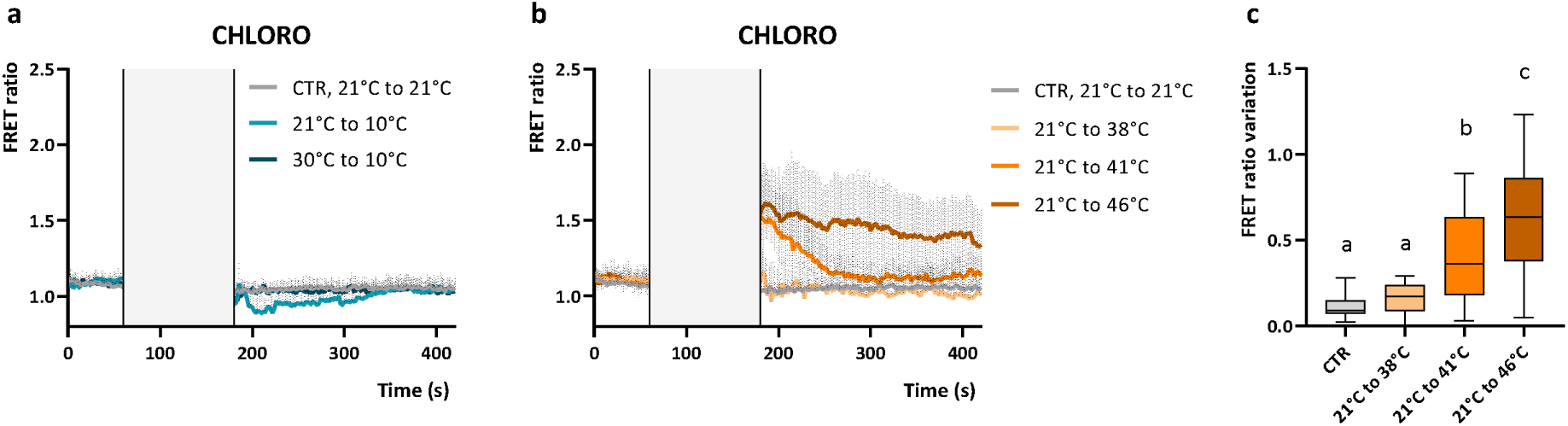
Chloroplast Ca^2+^ dynamics in *C. reinhardtii* cells in response to temperature stress. a, b. Averaged and normalized FRET Ratio ± SD of chloroplast YC3.6 in C. *reinhardtii* cells in response to cold shock (a, n > 11 cells) and heat stress (b, n > 17 cells). Grey rectangle indicate the treatment, 120 s of switch of perfusion at different temperature, 21°C-21°C (CTR), 21°C-10°C, 30°C-10°C, 21°C-38°C, 21°C-41°C, 21°C-46°C). c Maximum FRET Ratio variations triggered by heat stress respectively in the chloroplast (b, n > 17 cells). One-way ANOVA: P value < 0,05.

Heat or cold induced FRET Ratio change after stimulus of cytosol-localized YC3.6 probe was close to zero, indicating no alteration of the cytosolic [Ca^2+^] (Figure S4a, b). No *bona fide* FRET responses could be observed, as both fluorescent protein moieties signals (CFP and cpVenus) were similarly decreasing (Fig. S4d, e, f); this might be ascribable to temperature-dependent changes of YC3.6 fluorescence properties. The depletion of extracellular [Ca^2+^] did not alter the heat stress induced FRET ratio change (Figure S4c). Similarly, cells exposed to a cold shock (from 21°C (RT) to 10°C or from 30°C to 10°C) did not show chloroplast [Ca^2+^] elevations (Figure 4a). Neither the sensing of the absolute temperature of 10°C, nor an increased range of cooling (from 30°C to 10°C), triggered relevant changes in *C. reinhardtii* intracellular [Ca^2+^]. Sustained chloroplast-specific [Ca^2+^] elevations could be observed in cells exposed to 120 s of rapid increase of temperature from 21°C to 41 and 46°C (Figure 4b, c), characterized by *bona fide* FRET responses that last beyond the temperature stimulus (Figure S4g, h, i). Interestingly, a similar sustained [Ca^2+^] has been observed in *Arabidopsis* chloroplasts, when stimulated at 40°C (Lenzoni & Knight, 2019), suggesting the presence of a conserved mechanism at this level. No relevant cytosolic or chloroplast [Ca^2+^] elevations were observed in cells perfused with the same solutions after they had been equilibrated to RT (control, CTR), excluding the contribution of the act of switching between the perfusion solutions into the signalling responses.

### Exogenous applied bicarbonate and acetate induce large chloroplast [Ca^2+^] transients in photoautotrophic grown cells

*C. reinhardtii* CCM, and the involved bicarbonate (HCO_3_^-^) uptake system, are induced by limiting CO_2_ conditions. Different concentrations of limiting CO_2_ can lead to distinct acclimation strategies: ‘very low CO_2_’ (< 0.03% CO_2_), in which the cells predominantly concentrate Ci in the form of HCO_3_^−^ across the plasma membrane and chloroplast envelope, and ‘low CO_2_’ (0.03-0.5% CO_2_), when the cell switches to a CO_2_-based uptake system (Atkinson et al., 2016; H. Gao, Wang, Fei, Wright, & Spalding, 2015; Mackinder, 2018; Y. Wang & Spalding, 2014). To experimentally address the role of intracellular Ca^2+^ signalling in *C. reinhardtii* carbon sensing and carbon concentrating mechanisms, cytosolic and chloroplast [Ca^2+^] were monitored in YC3.6 expressing lines grown in CO_2_-limiting photoautotrophic conditions. In particular, cells were grown in shaking flasks in minimal medium (TP), without air bubbling. Sodium bicarbonate (NaHCO_3_) or acetate were dissolved in TP medium at the indicated concentrations and rapidly perfused to resupply different carbon sources to the cells. The typical acetate concentration of TAP medium was applied (17 mM), whereas different NaHCO_3_ concentrations were tested (5-40 mM). To avoid the generation of acid-induced cytosolic [Ca^2+^] elevations, previously reported in *C. reinhardtii* to induce deflagellation (Glen L. Wheeler, Joint, & Brownlee, 2007), the pH of all the perfused solutions was adjusted to 7.

NaHCO_3_ and acetate perfusion did not lead to large cytosolic [Ca^2+^] elevations (defined as > 0.2-point variation of the maximal FRET ratio value, Figure 5a, b), with only 17.8 % and 30 % of the cells displaying a maximal FRET ratio change above 0.2. Nonetheless, a small but significant increase in the mean maximal FRET ratio was observed in both treatments (Figure 5c): FRET ratio signal rapidly returns to resting values when cells were treated with acetate, whereas it shows a much slower decay when triggered by 40 mM NaHCO_3_ (Figure 5a, b). The maximum FRET ratio variation is comparable between the two treatments (Figure 5c, left panel), but the “Area Under the Curve” (AUC, indicating both the amplitude and the duration of the Ca^2+^ transient) is indeed significantly higher in the bicarbonate treated cells (Figure 5c, right panel), suggesting a longer duration of the Ca^2+^ transient.

**Figure 5.**
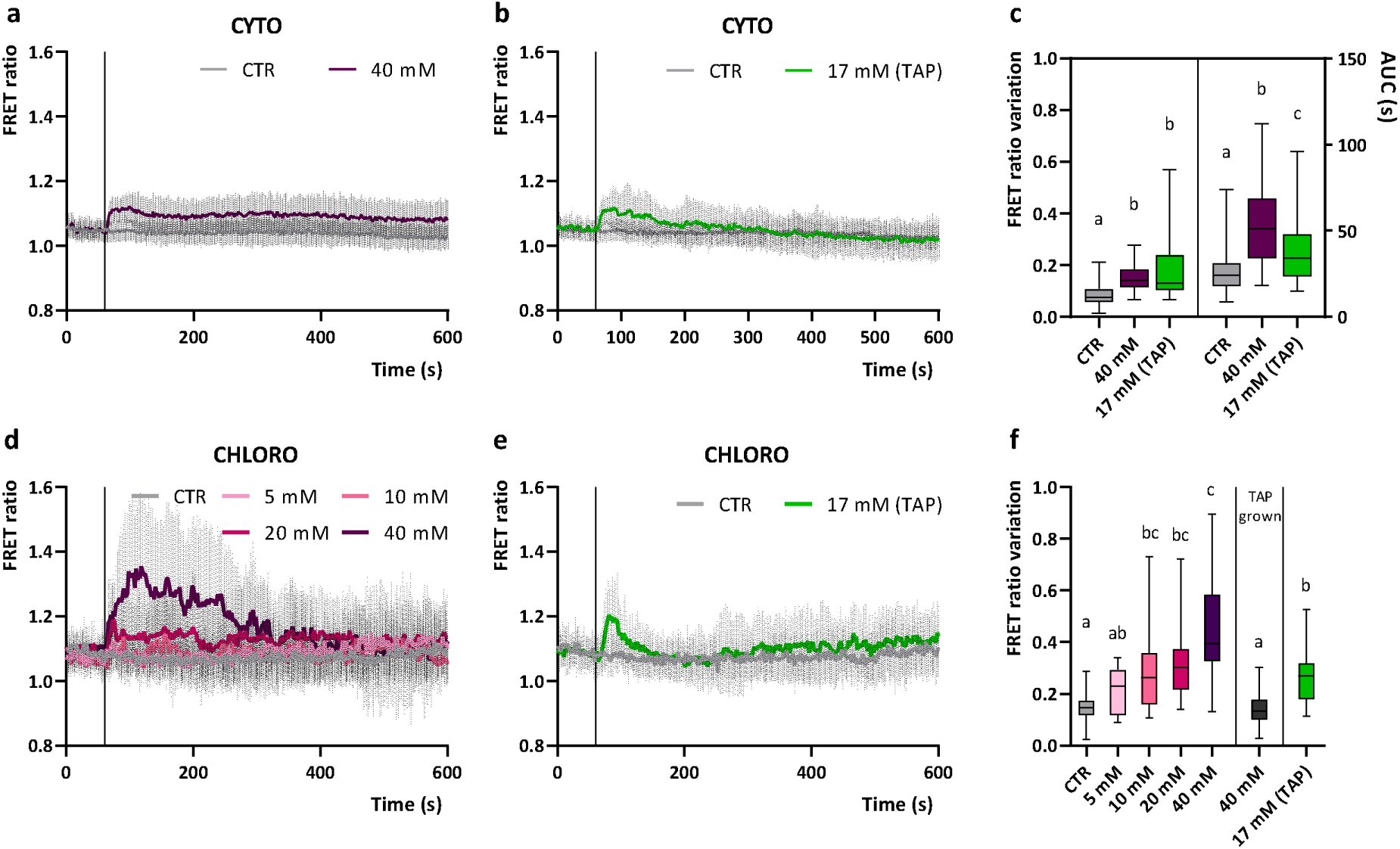
Cytosolic and chloroplast Ca^2+^ dynamics in *C. reinhardtii* cells in the response to different exogenous Ci sources. a, b. Averaged and normalized FRET Ratio ± SD of cytosolic YC3.6 in C. *reinhardtii* cells in response to sodium bicarbonate (a, n > 45 cells) or acetate (b, n > 40 cells) at the indicated concentration in TP medium, pH 7 (black vertical line indicates the onset of the stimulus, 60 s). d, e. Averaged and normalized FRET Ratio ± SD of chloroplast YC3.6 in C. *reinhardtii* cells in response to sodium bicarbonate (d, n > 10 cells) or acetate (e, n > 23 cells) at the indicated concentrations in TP medium, pH 7 (black vertical line indicates the onset of the stimulus, 60 s). c, f Maximum FRET Ratio variations triggered by sodium bicarbonate or acetate respectively in cytosol (c, n > 40 cells) and in the chloroplast (f, n > 10 cells). One-way ANOVA: P value < 0,05.

Both NaHCO_3_ and acetate perfusions also induced chloroplast-specific [Ca^2+^] elevations, even though with different kinetic properties. To further investigate the chloroplast sensitivity to exogenous Ci, the dose dependency of the response to different concentrations of NaHCO_3_ (5-40 mM) was evaluated (Figure 5d). The chloroplast maximal FRET ratio variation showed a moderate correlation with the intensity of the stimulus applied, obtaining a significant response with 10 mM and 20 mM NaHCO_3,_ and a maximal amplitude of it with 40 mM NaHCO_3_ treatment (Figure 5f). It is clear also that the proportion of cells exhibiting chloroplast [Ca^2+^] elevations change with increasing sodium bicarbonate concentrations (Figure S6): for instance, 40 mM NaHCO_3_ resulted in a FRET ratio elevation above 0.2-points in over 92% of the observed cells. The spatial characteristics of the sodium bicarbonate-induced chloroplast [Ca^2+^] elevations were also examined. The triggered [Ca^2+^] elevations were observed across all chloroplast regions, not solely in the pyrenoid (Figure S5c). However, the pyrenoid resting steady-state FRET ratio was significantly higher within the pyrenoid relative to the rest of the chloroplast in photoautotrophic grown cells, similar to our observations in mixotrophic grown cells (Figure S5b).

In the case of mixotrophically grown cells (TAP medium), 40 mM sodium bicarbonate did not induce any significant chloroplast [Ca^2+^] elevation (Figure 5f). Acetate in TAP medium, the commonly used reduced carbon source for *C. reinhardtii* mixotrophic growth, can indeed suppress CCM induction (Fett & Coleman, 1994; Moroney, Kitayama, Togasaki, & Tolbert, 1987). The acetate inhibition of this Ca^2+^-mediated response might thus suggest the presence of a link between the CCM functioning, external carbon sensing mechanisms and intracellular Ca^2+^ signalling. Acetate itself triggered a chloroplast-specific single rapid [Ca^2+^] elevation, significantly higher than control (CTR) (Figure 5e), indicating a further connection between intracellular Ca^2+^ signalling and external carbon sensing.

Cells grown in autotrophic conditions, but daily supplemented with 20 mM or 40 mM sodium bicarbonate as exogenous carbon source, do not show significant chloroplast Ca^2+^ transients in response to 40 mM sodium bicarbonate resupply (Figure 6). The daily supplementation of sodium bicarbonate to autotrophically grown cultures may prevent activation of the CCM, which is induced by low CO_2_ concentrations (Mackinder, 2018; Ruiz-Sola et al., 2023). Accordingly, we observed slightly lower levels of CARBONIC ANHYDRASE 3 (CAH3) and of the large subunit of RuBisCO (RbcL) in the sodium bicarbonate supplemented cultures, compared to the control (Figure S7), although these were not significant (Polukhina, Fristedt, Dinc, Cardol, & Croce, 2016) (Figure S7).

**Figure 6.**
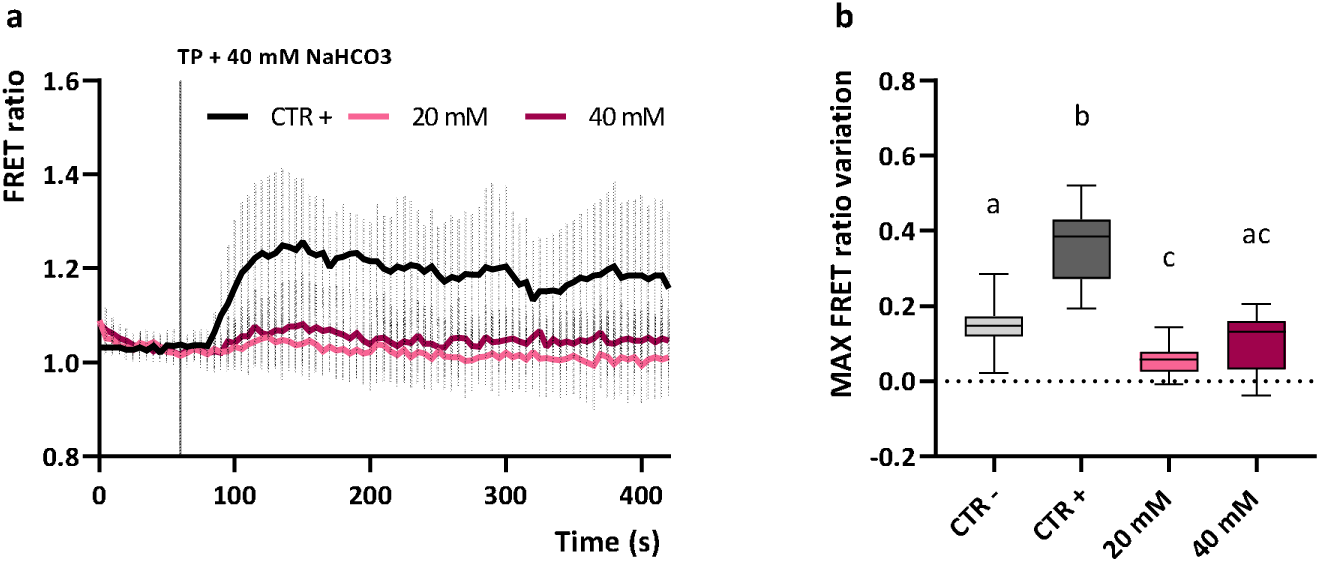
Chloroplast Ca^2+^ dynamics in *C. reinhardtii* cells in response to 40 mM sodium bicarbonate (TP + 40 mM NaHCO_3_). a. Averaged and normalized FRET Ratio ± SD of chloroplast YC3.6 in C. *reinhardtii* cells in response to 40 mM sodium bicarbonate (n > 10 cells) in autotrophic grown cultures (CTR +) and in daily supplemented cultures with sodium bicarbonate at the indicated concentration in TP medium, pH 7 (black vertical line indicates the onset of the stimulus, 60 s). b. Maximum FRET Ratio variations triggered by sodium bicarbonate in the chloroplast (n > 10 cells) in the different autotrophic grown cultures reported in a. CTR - represent values from untreated cells grown in comparable conditions and reported also in Figure 5f as CTR. One-way ANOVA: **, P value < 0,001.

Taken together these data showed that photoautotrophically grown *C. reinhardtii* cells are sensitive to different external Ci sources and that cytosolic and chloroplast specific Ca^2+^ signalling might be involved in their sensing mechanisms.

## DISCUSSION

The mechanisms of perception of extracellular stresses and their subsequent transduction into physiological responses are required by nearly all eukaryotic cells to adapt to the environment. The requirement for Ca^2+^-dependent signalling pathways in many of these environmental responses has been shown to be conserved among algae, plants, or even animal cells, displaying common elements in the underlying cellular mechanisms (M. Pivato et al., 2023; Stael et al., 2012). Here, intracellular Ca^2+^ signalling is demonstrated to be likely involved in the responses of green algae to several different environmental stressors, including osmotic stimuli, temperature fluctuations and Ci availability (Table 1).

**Table 1.** Summary of the main cytosolic and chloroplast Ca^2+^ signalling responses detected in *C. reinhardtii* cells in response to abiotic stresses.

|  |  | Cytosol |  |  | Chloroplast stroma |  |  |
| --- | --- | --- | --- | --- | --- | --- | --- |
| | | $\text{Ca}^{2+}$ response | Description | Reference | $\text{Ca}^{2+}$ response | Description | Reference |
| High light | Blue light | ✗ | No $\text{Ca}^{2+}$ transient observed | Pivato *et al.*, 2023 | ✓ | Chloroplast-specific $[\text{Ca}^{2+}]$ elevation | Pivato *et al.*, 2023 |
| | Red light | ✗ | No $\text{Ca}^{2+}$ transient observed | Pivato *et al.*, 2023 | ✓ | Chloroplast-specific $[\text{Ca}^{2+}]$ elevation | Pivato *et al.*, 2023 |
| ROS | $\text{H}_2\text{O}_2$ | N/A | Not assessed | / | ✓ | 1 mM, induced $[\text{Ca}^{2+}]$ elevation | Pivato *et al.*, 2023 |
| Nutrient resupply | $\text{NH}_4^+$ | ✗ | No $\text{Ca}^{2+}$ transient observed | This work | ✗ | No $\text{Ca}^{2+}$ transient observed | This work |
| | $\text{PO}_4^{2-}$ | ✗ | No $\text{Ca}^{2+}$ transient observed | This work | ✗ | No $\text{Ca}^{2+}$ transient observed | This work |
| Hyper-osmotic | NaCl | ✓ | 100, 200 and 300 mM induced $[\text{Ca}^{2+}]$ elevation | This work, Bickerton *et al.*, 2016 | ✓ | 200 and 300 mM induced $[\text{Ca}^{2+}]$ elevation | This work |
| | Sorbitol | ✓ | 360 mM induced $[\text{Ca}^{2+}]$ elevation | This work | ✓ | 360 mM induced $[\text{Ca}^{2+}]$ elevation | This work |
| Hypo-osmotic | milliQ $\text{H}_2\text{O}$ | ✓ | Induced $[\text{Ca}^{2+}]$ spikes with higher osmolarity adaptation | This work, Bickerton *et al.*, 2016 | ✗* | No $\text{Ca}^{2+}$ signalling observed during stimulus *only after, hyper-osmotic | This work |
| Low temperature | $10^\circ\text{C}$ | ✗ | No $\text{Ca}^{2+}$ transient observed | This work | ✗ | No $\text{Ca}^{2+}$ transient observed | This work |
| High temperature | 38, 41, $46^\circ\text{C}$ | ✗ | No $\text{Ca}^{2+}$ transient observed | This work | ✓ | 41 and $46^\circ\text{C}$ chloroplast-specific $[\text{Ca}^{2+}]$ sustained elevation | This work |
| Bicarbonate | $\text{NaHCO}_3$ | ✓* | * Observed small, but significant, 40 mM induced $[\text{Ca}^{2+}]$ elevations, (see "Discussion") | This work | ✓ | 10, 20 and 40 mM induced $[\text{Ca}^{2+}]$ elevations | This work |
| Acetate | $\text{CH}_3\text{COOH}$ | ✓* | * Observed small, but significant, 17 mM induced $[\text{Ca}^{2+}]$ elevations, (see "Discussion") | This work | ✓ | 17 mM induced $[\text{Ca}^{2+}]$ peak | This work |

**Table 1. Summary of the main cytosolic and chloroplast $\text{Ca}^{2+}$ signalling responses detected in *C. reinhardtii* cells in response to abiotic stresses.**
|  |  | Cytosol |  |  | Chloroplast stroma |  |  |
| --- | --- | --- | --- | --- | --- | --- | --- |
| | | $\text{Ca}^{2+}$ response | Description | Reference | $\text{Ca}^{2+}$ response | Description | Reference |
| High light | Blue light | ✗ | No $\text{Ca}^{2+}$ transient observed | Pivato *et al.*, 2023 | ✓ | Chloroplast-specific $[\text{Ca}^{2+}]$ elevation | Pivato *et al.*, 2023 |
| | Red light | ✗ | No $\text{Ca}^{2+}$ transient observed | Pivato *et al.*, 2023 | ✓ | Chloroplast-specific $[\text{Ca}^{2+}]$ elevation | Pivato *et al.*, 2023 |
| ROS | $\text{H}_2\text{O}_2$ | N/A | Not assessed | / | ✓ | 1 mM, induced $[\text{Ca}^{2+}]$ elevation | Pivato *et al.*, 2023 |
| Nutrient resupply | $\text{NH}_4^+$ | ✗ | No $\text{Ca}^{2+}$ transient observed | This work | ✗ | No $\text{Ca}^{2+}$ transient observed | This work |
| | $\text{PO}_4^{2-}$ | ✗ | No $\text{Ca}^{2+}$ transient observed | This work | ✗ | No $\text{Ca}^{2+}$ transient observed | This work |
| Hyper-osmotic | NaCl | ✓ | 100, 200 and 300 mM induced $[\text{Ca}^{2+}]$ elevation | This work, Bickerton *et al.*, 2016 | ✓ | 200 and 300 mM induced $[\text{Ca}^{2+}]$ elevation | This work |
| | Sorbitol | ✓ | 360 mM induced $[\text{Ca}^{2+}]$ elevation | This work | ✓ | 360 mM induced $[\text{Ca}^{2+}]$ elevation | This work |
| Hypo-osmotic | milliQ $\text{H}_2\text{O}$ | ✓ | Induced $[\text{Ca}^{2+}]$ spikes with higher osmolarity adaptation | This work, Bickerton *et al.*, 2016 | ✗* | No $\text{Ca}^{2+}$ signalling observed during stimulus *only after, hyper-osmotic | This work |
| Low temperature | 10°C | ✗ | No $\text{Ca}^{2+}$ transient observed | This work | ✗ | No $\text{Ca}^{2+}$ transient observed | This work |
| High temperature | 38, 41, 46 °C | ✗ | No $\text{Ca}^{2+}$ transient observed | This work | ✓ | 41 and 46°C chloroplast-specific $[\text{Ca}^{2+}]$ sustained elevation | This work |
| Bicarbonate | $\text{NaHCO}_3$ | ✓* | * Observed small, but significant, 40 mM induced $[\text{Ca}^{2+}]$ elevations, (see "Discussion") | This work | ✓ | 10, 20 and 40 mM induced $[\text{Ca}^{2+}]$ elevations | This work |
| Acetate | $\text{CH}_3\text{COOH}$ | ✓* | * Observed small, but significant, 17 mM induced $[\text{Ca}^{2+}]$ elevations, (see "Discussion") | This work | ✓ | 17 mM induced $[\text{Ca}^{2+}]$ peak | This work |

### Nutrient resupply does not affect intracellular [Ca^2+^]

The importance of Ca^2+^ signalling in nutrient sensing is rapidly emerging from recent studies: nitrate Ca^2+^ signalling is essential for N sensing in plants (Riveras et al., 2015), but not in diatoms, where Ca^2+^ signalling is employed for phosphate sensing rather than nitrate (Helliwell et al., 2021). Conversely, the results herein presented show that in *C. reinhardtii* cells intracellular Ca^2+^ signalling is not involved in exogenous ammonium or phosphate sensing (Figure 1). Nitrate and ammonium are the primary N sources for many organisms, including *C. reinhardtii*. Ammonium is preferentially assimilated from these organisms, since its integration into organic form is energetically favourable; however, nitrate wider distribution and higher concentration in natural environments makes it the much more available N source (Fernandez & Galvan, 2007; Sanz-Luque, Chamizo-Ampudia, Llamas, Galvan, & Fernandez, 2015). YC3.6 expressing lines herein adopted were generated from a UVM4 background strain (Neupert et al., 2009), which carry the *nit1* and *nit2* mutations, and cannot grow on nitrate as the sole N source. Thus, from these data the presence in *C. reinhardtii* of a conserved nitrate-dependent Ca^2+^ signalling mechanism for N sensing cannot be excluded. Future experiments will be indeed addressed to test this hypothesis in newly obtained YC3.6 expressing lines that can efficiently assimilate nitrate.

### Intracellular [Ca^2+^] elevations in response to hyper-osmotic shock

The results herein reported suggest that *C. reinhardtii* intracellular Ca^2+^ signalling is likely involved in the sensing mechanisms of several other environmental factors. Hyper-osmotic induced cytosolic [Ca^2+^] elevations were observed, whose amplitude were dependent of the strength of the stimulus, with larger Ca^2+^ elevations observed at the higher NaCl concentrations (Figure 2a). The observed transients differed from the ones previously reported with comparable stimuli (Bickerton et al., 2016), which showed very rapid spiking-shaped kinetics. However, distinct Ca^2+^ signalling responses of walled and wall-less strains to osmotic stimuli have already been reported, highlighting a moderate variability in the hyper-osmotic induced Ca^2+^ signalling of different *C. reinhardtii* strains. Similarly, cell-wall deficient YC3.6 expressing lines (UVM4 background, (Neupert et al., 2009)) did not display any significant cytosolic [Ca^2+^] elevation in response to hypo-osmotic stimuli (a, b, c) but reported comparable hypo-osmotic induced cytosolic [Ca^2+^] elevations when acclimated to a higher osmolarity (TAP + 100 mM sorbitol) (2a, b, c). These findings further support the previously advanced hypothesis, for which cell wall may contribute to osmotic signalling; whether this effect is the physiological consequence of its absence/presence or the product of its interaction with the plasma membrane is not yet clear. In addition, the findings herein reported reveal previously unreported hyper-osmotic induced chloroplast [Ca^2+^] elevations (Figure 2d), observed only above a threshold stimulus (200 mM NaCl), required to trigger the response. Some aspects of these Ca^2+^ elevations are comparable to those observed in vascular plants, where studies using the Ca^2+^-responsive bioluminescent reporter aequorin revealed in *Arabidopsis* both cytosolic and chloroplast NaCl-induced [Ca^2+^] elevations, even though with different spatial and temporal properties (Sello et al., 2018; Tracy, Gilliham, Dodd, Webb, & Tester, 2008). Interestingly, both the cytosolic and chloroplast hyper-osmotic responses are likely to be caused by an osmotic effect of the treatment, since iso-osmotic sorbitol treatments induced comparable compartment-specific [Ca^2+^] increases (Figure 2e, f). The chloroplast volume alterations following the hyper-osmotic shocks (Figure 3a, d), together with the transient increase of intracellular [Ca^2+^], might indicate the activation of mechanosensitive and/or osmosensitive ion channels at the level of the plasma membrane, chloroplast envelope, thylakoid membranes, or even other subcellular compartments. Indeed, the depletion of extracellular [Ca^2+^] did not alter the hyper-osmotic-induced Ca^2+^ signature in both the subcellular compartments (Figure 2e, f), suggesting a relevant role of intracellular compartments as main Ca^2+^ sources in these responses. Examples of candidate molecular players could be represented by members of the transient receptor potential (TRP) subfamily of cation channels, composed by at least 21 members in *C. reinhardtii* (Arias-Darraz et al., 2015), and activated by osmotic stimuli, the PLASTID ENVELOPE ION CHANNELS (PECs), found in *A. thaliana* to be involved in stress-triggered stromal Ca^2+^ release (Völkner et al., 2021), or by mechanosensitive ion channels, with seven members in *C. reinhardtii* (MSC1-MSC7) (Edel & Kudla, 2015).

### Heat stress induced chloroplast [Ca^2+^] elevations

Temperature fluctuations are environmental stressors that severely affect growth and productivity in plants and algae. The mechanisms for temperature sensing however, specifically the early sensing and signalling events, are still an open research topic in these organisms, even if Ca^2+^ signalling has been reported to be frequently involved. Heat stress has already been reported to trigger elevations of cytosolic and nuclear H_2_O_2_ levels in *C. reinhardtii* cells exposed to 40°C for 30 min, without major detrimental effects (Niemeyer et al., 2021). Moreover, several recent studies reported wide differences in the physiological responses of cells adapted to moderate (35°C) or acute (40°C) high temperatures (Mattoon et al., 2023; N. Zhang et al., 2022). In diatoms transient cytosolic [Ca^2+^] elevations are triggered in response to rapid cooling, similarly to what has been already observed in plant and animal cells (Kleiner et al., 2022; H. Knight et al., 1996; M. R. Knight & Knight, 2012; Q. Liu et al., 2021). In *C. reinhardtii* cells no relevant cytosolic or chloroplast [Ca^2+^] elevations could be detected in response to cold shocks (from 21°C (RT) to 10°C or from 30°C to 10°C). The equilibration of the small volume of cold perfusate with room temperature prevented from testing lower temperatures, opening to the possibility that a stronger temperature decreasing, or lower absolute temperatures, could be required to trigger any intracellular [Ca^2+^] elevation in *C. reinhardtii* cells, similar to the case of diatoms and plants. Heat stress trigger transient elevation of the [Ca^2+^] in the plant cell cytosol, in which the influx of extracellular calcium is crucial, as demonstrated upon application of calcium chelators or calcium channels blockers (F. Gao et al., 2012; Saidi et al., 2009; Wu, Luo, Vignols, & Jinn, 2012). Here, no heat stress induced FRET ratio change was detected in the cytosol (Figure S4b) of *C. reinhardtii* cells that persisted after the removal of the temperature stimulus. Differently, significant chloroplast-specific [Ca^2+^] elevations were evident in cells exposed to rapid increases of temperature from 21°C to 41 and 46°C (Figure 4b), as previously reported for plant chloroplasts exposed to heat stress (Lenzoni & Knight, 2019). The reported responses are characterized by *bona fide* FRET ratio changes and displayed characteristic kinetics, independent from the temperature stimulus dynamics (Figure S4).

Temperature can shape chloroplast physiology (Mathur, Agrawal, & Jajoo, 2014), influencing the catalytic activity of different enzymes involved in the carbon fixation reactions (Sage, Way, & Kubien, 2008) or in the fatty acid biosynthesis, but also altering membranes fluidity and affecting plastoquinone diffusion, ultimately affecting photosynthetic electron transport (Pshybytko, Kruk, Kabashnikova, & Strzalka, 2008). Moreover, temperature elevations can have significant effects on functional chloroplast proteins, for instance inhibiting photosynthetic complexes activity (Murata, Takahashi, Nishiyama, & Allakhverdiev, 2007; Ueno et al., 2016). Together, these temperature-related effects might account for the high chloroplast sensitivity to temperature fluctuations and may represent the base for the evolution of its key role in the thermotolerance response (Dickinson et al., 2018). Increasing evidence is indeed reporting in land plants a crucial role of chloroplast Ca^2+^ signalling as an intracellular second messenger in this process, even though the underlying molecular mechanisms have not yet been clarified (Lenzoni & Knight, 2019; Pollastri, Sukiran, Jacobs, & Knight, 2021).

Heat stress-induced [Ca^2+^] elevation in plant cells was suggested to be mediated by the opening of specific calcium-permeable channels as a consequence of increased membrane fluidity at elevated temperatures (Saidi et al., 2009). Good candidates for calcium-permeable channels are cyclic nucleotide-gated channels (CNGC), non-selective inward cation channels containing domains for the binding of cyclic nucleotides and calmodulin, previously shown to be involved in *Arabidopsis* heat stress-induced [Ca^2+^] elevations (F. Gao et al., 2012). The *C. reinhardtii* genome also contains three CNGC genes, that could have a role in intracellular Ca^2+^ dynamics following heat stress, as well as their plant counterpart (Edel & Kudla, 2015; Matteo Pivato & Ballottari, 2021); nevertheless, no CNGC have been found in the chloroplast yet and their functions and subcellular localization are still unknown.

### The role of intracellular Ca^2+^ signalling in exogenous Ci sensing

This study revealed an interesting link between intracellular Ca^2+^ signalling and exogenous supply of different Ci sources, NaHCO_3_ and acetate. A key role of Ca^2+^ in the CO_2_ sensing mechanism and CCM of *C. reinhardtii* has already been reported (L. Wang et al., 2016), but current knowledge about the first steps of the sensing and intracellular signalling mechanism is still fragmented. Here, cytosolic and chloroplast [Ca^2+^] dynamics were measured upon NaHCO_3_ or acetate perfusion in photoautotrophically grown and CO_2_ limited cells (Figure 5). A recent work using an aequorin-based imaging system described HCO_3_^−^-induced cytosolic [Ca^2+^] increases in *Arabidopsis* leaves and guard cells (Tang et al., 2020) but, to our knowledge, comparable responses have not been reported in chloroplasts so far, nor in algal systems. The amplitude and kinetic properties of these responses differed for the applied stimulus, showing also a moderate dose dependency in the case of NaHCO_3_ induced chloroplast [Ca^2+^] transients. Both 40 mM NaHCO_3_ and 17 mM acetate might have a mild hyperosmotic effect on *C. reinhardtii* cells; however, as 100 mM NaCl treatment did not influence chloroplast [Ca^2+^] (Figure 2c) it can be ruled out that the NaHCO_3_ and acetate induced chloroplast Ca^2+^ responses are caused by a hyper-osmotic effect.

Large cytosolic [Ca^2+^] elevations in response to NaHCO_3_ or acetate were not detected even though both treatments significantly perturbed the basal steady-state FRET ratio (Figure 5a-c). Since 100 mM NaCl stimulus induces significant transient cytosolic [Ca^2+^] elevation (Figure 2c, e), it must be considered that 40 mM NaHCO_3_ treatment might as well have a mild hyper-osmotic effect, as it causes a significant shrinking of the cell area (Figure S8). Nevertheless, NaHCO_3_ induced cytosolic [Ca^2+^] elevation showed unique rapid induction and longer duration characteristics, that differed from the NaCl induced Ca^2+^ response. Furthermore, also a minor pH effect of NaHCO_3_ and acetate stimuli cannot totally be excluded, since both could cross the plasma membrane respectively as CO_2_ and undissociated acetate (at a pH below acetic acid p*K*_a_ 4.75) (Casal, Cardoso, & Leao, 1996; Casal, Paiva, Queirós, & Soares-Silva, 2008), resulting in cytosolic acidification and acid-induced cytosolic [Ca^2+^] elevations (G. L. Wheeler, Joint, & Brownlee, 2008). Nonetheless, the pH of the TP + 40 mM NaHCO_3_ and TAP solutions was adjusted to 7 prior stimulation, widely mitigating their potential acidification effect. However, further characterization of the NaHCO_3_ or acetate cytosolic Ca^2+^ response is needed to reliably exclude potential osmotic and pH effects on the cytosolic [Ca^2+^].

Distinct Ca^2+^ signalling toolkits could account for the observed [Ca^2+^] elevations, as each organelle can display a unique set of Ca^2+^ related molecular players. The transient receptor potential (TRP) Ca^2+^-channel TRP2, putatively located in the chloroplast envelope, have been proposed to relay Ca^2+^ signals from the chloroplast to the cytosol (Christensen, Dave, Mukherjee, Moroney, & Machingura, 2020), even though experimental evidence in this direction is still missing. In CO_2_-limiting conditions, significant increases of [Ca^2+^], reported through a chemical Ca^2+^ dye (Calcium Green-1, AM), have been observed in the pyrenoid region of the chloroplast, along with a CAS association with tubule-like structures of it (L. Wang et al., 2016). Moreover, the perturbation of intracellular Ca^2+^ homeostasis by a Ca^2+^ chelator or calmodulin antagonists impair the accumulation of two specific HCO_3_^-^ transporters, high-light activated 3 (HLA3) and low-CO_2_–inducible gene A (LCIA), both essential for the HCO_3_^-^ uptake into the chloroplast stroma (L. Wang et al., 2016). Therefore, from the data herein reported, a model could be envisaged in which Ci induced cytosolic and chloroplast [Ca^2+^] elevations play a role in modulating the response to the availability of different external Ci sources. SLAC1 anion channel has been proposed as one of the physiologically relevant CO_2_/HCO_3_^−^ sensors in plant guard cells, regulating stomatal movements (J. Zhang et al., 2018). Putative homologs could play a similar role in microalgae, even though they have not been identified so far; however, the presence of other HCO_3_^-^ or acetate-regulated ion channels that could mediate a comparable function in *C. reinhardtii* could not be excluded. Even so, the findings herein presented, report the involvement of chloroplast Ca^2+^ signalling in the sensing mechanism of exogenous acetate and HCO_3_^-^ in green algae, helping to elucidate the so far incomplete picture on the first steps of it.

Overall, the results presented in this work report abiotic stress-induced stimulus-specific Ca^2+^ transients in *C. reinhardtii*, suggesting the potential involvement of intracellular Ca^2+^ signalling in the perception mechanisms of different environmental stimuli in green algae. The use of the previously obtained YC3.6 expressing lines allowed us to monitor cytosolic and chloroplast [Ca^2+^] independently, helping to identify unique [Ca^2+^] transients that might trigger specific downstream responses. In summary, cytosol-specific [Ca^2+^] transients were observed in response to hypo-osmotic stress, suggesting that the expansion of cell volume and the risk of bursting primarily affects cytosolic signalling. Conversely, cell contraction caused by hyper-osmotic stress may have an impact on both compartments, triggering characteristic cytosolic and chloroplast [Ca^2+^] elevations. Exogenous supply of different Ci sources and heat stress primarily affects chloroplast [Ca^2+^], suggesting a major role of the chloroplast in the perception of these environmental stimuli. As both light and Ci sources preferentially influence chloroplast bioenergetic processes, it is reasonable to expect that this subcellular compartment could primarily and autonomously respond to high light stress, H_2_O_2_, photo-oxidative stress (M. Pivato et al., 2023) and/or Ci availability. The central role of Ca^2+^ in these sensing mechanisms appears to be conserved with land plants for a major part of the obtained responses, even if with some differences: nitrate resupply and cold stress trigger [Ca^2+^] transients in land plants, but not in *C. reinhardtii*. These differences might be ascribable to a combination of distinct Ca^2+^ signalling toolkits found in plants and green algae and even more extended divergencies in stress response and cell morphological and physiological characteristics. Elucidating the molecular basis of these differences will provide new understandings of the mechanisms that microalgae exploit to respond to specific natural conditions and will also provide new insights into the evolution of eukaryotic fundamental cellular processes.

## ACKNOWLEDGEMENTS

This work was supported by the research program “Dipartimento di Eccellenza 2018-2022” (Ministero dell’Università e della Ricerca, DIPCEL5) (to M.P. and M.B.), by Cariverona Foundation (research grant CARIVERONA-HABITAT-2022 to M.B.) and by “EMBO Scientific Exchange Grant” (EMBO) (to M.P.).

