## Supplementary material for "Abiotic stress-induced chloroplast and cytosolic Ca^2+^dynamics in the green alga *Chlamydomonas reinhardtii*": Figure S

### SUPPORTING INFORMATION

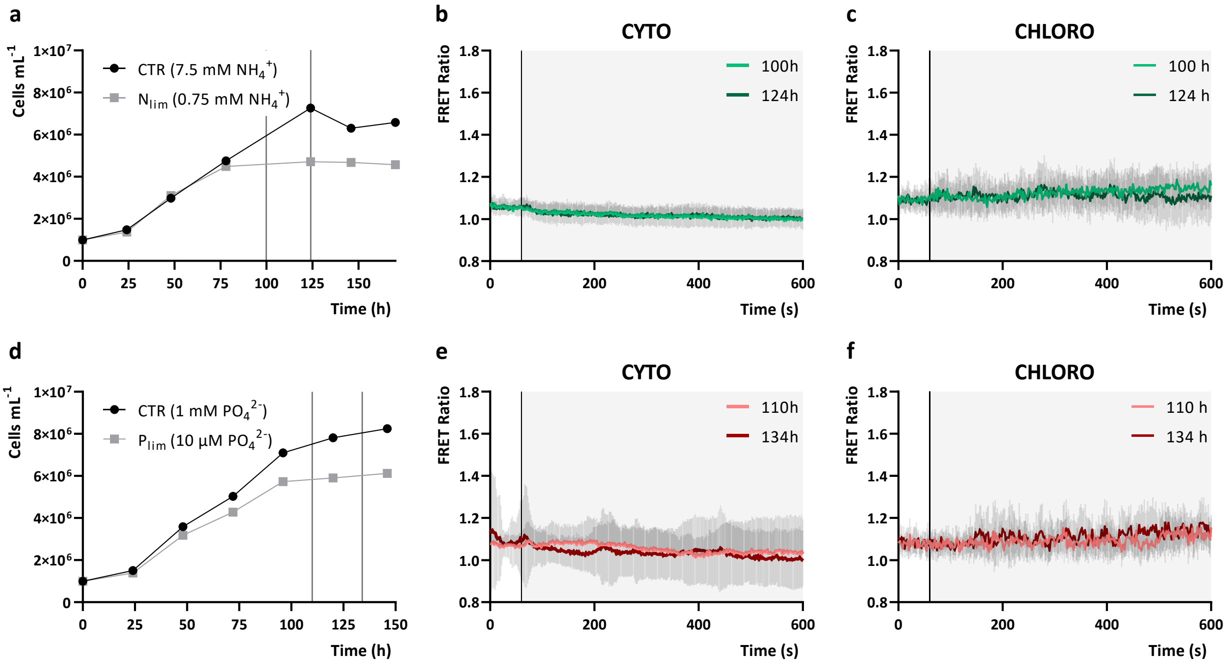


**Figure S1. Increases in environmental ammonium or phosphates doesn't affect cytosolic and chloroplast stroma $Ca^{2+}$ transients in photoautotrophic grown cells.** a, d. Photoautotrophic growth over time of *C. reinhardtii* UVM4 cells in standard TP medium with ammonium- (a) or phosphate-replete (d) (CTR, 7.5 mM $NH_4^+$ and 1 mM $PO_4^{2-}$ respectively), versus ammonium- or phosphate-limited ($N_{lim}$ 0.75 mM $NH_4^+$ and $P_{lim}$ 10 µM $PO_4^{2-}$ respectively) conditions (n = 1). Black vertical lines in the graphs indicate the time-points of the growth curve at which nutrient deplete YC3.6 expressing cells were harvested to perform nutrient resupply experiments (reported in different colour shades in b, c, e and f). b, c. Averaged and normalized FRET Ratio ± SD of cytosolic (b, n > 53 cells) and chloroplast (c, n > 13 cells) YC3.6 in *C. reinhardtii* cells in response to ammonium resupply (TAP, 7.5 mM $NH_4^+$) (black vertical line indicate the onset of the stimulus, at 60 s). e, f. Averaged and normalized FRET Ratio + SD of cytosolic (e, n > 59 cells) and chloroplast (f, n > 9 cells) YC3.6 in *C. reinhardtii* cells in response to phosphate resupply (TAP, 1 mM $PO_4^{2-}$) (black vertical line indicate the onset of the stimulus, at 60 s).

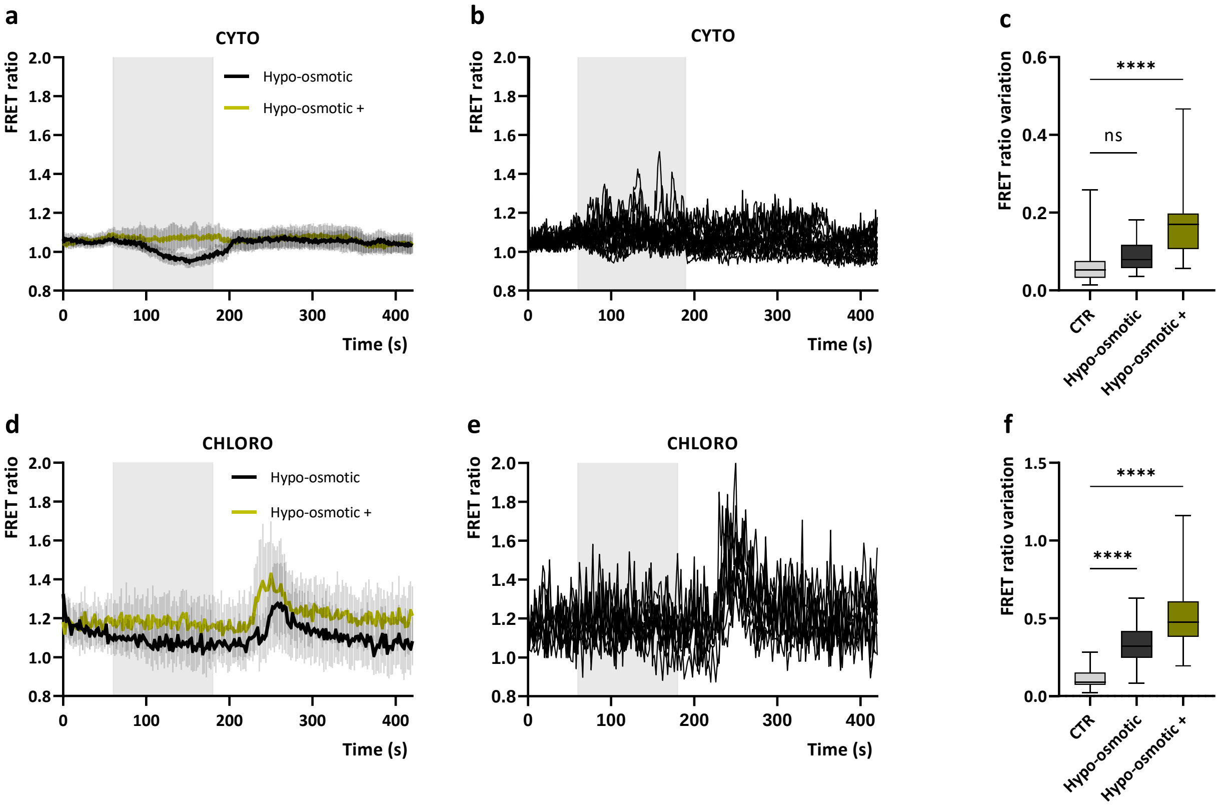


**Figure S2. Cytosolic and chloroplast $Ca^{2+}$ dynamics in *C. reinhardtii* cells in response to hypo-osmotic stimuli.** a, d. Averaged and normalized FRET Ratio ± SD of cytosolic (a, n > 44 cells) and chloroplast (b, n > 21 cells) YC3.6 in C. *reinhardtii* cells in response to hypo-osmotic stimuli (grey rectangle indicate the treatment, 120 s of switch from growth medium (TAP) to deionized water + 0.34 mM $CaCl_2$, black line "Hypo-osmotic", or from growth medium (TAP) + 100 mM Sorbitol to deionized water + 0.34 mM $CaCl_2$, yellow line "Hypo-osmotic +"). b, e. Representative FRET Ratio single traces of cytosolic (b, n = 14 cells) and chloroplast (e, n = 10 cells) YC3.6 in C. *reinhardtii* cells acclimated to TAP + 100 mM Sorbitol (selected cells with a maximal FRET Ratio variation $\geq$ 0.2) in response to hypo-osmotic stimuli (grey rectangle indicate the treatment, 120 s of switch to deionized water + 0.34 mM $CaCl_2$). c, f. Maximum FRET Ratio variations of cytosolic (c, n > 44 cells) and chloroplast (f, n > 21 cells) YC3.6 in C. *reinhardtii* cells in response to hypo-osmotic shocks (referred to a and d respectively), compared to control (no stimulus, CTR). One-way ANOVA: **** = P value < 0.0001.

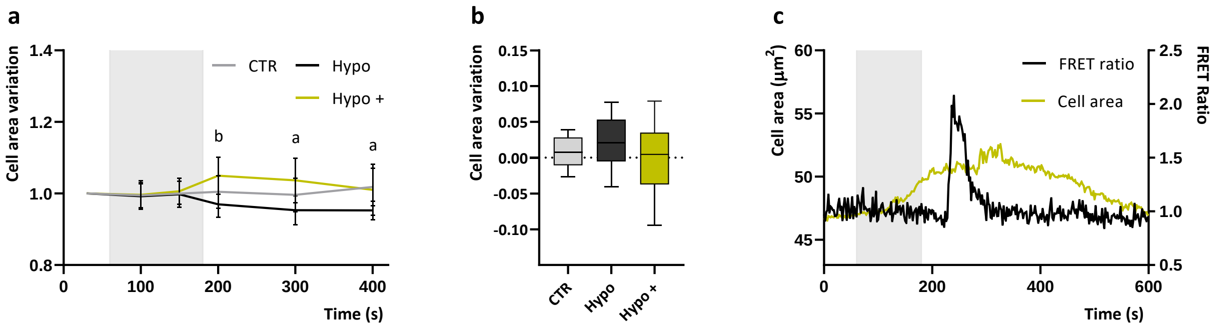


**Figure S3. *C. reinhardtii* cell area variation in response to hypo-osmotic stimuli.** a. Averaged and normalized cell area measurements ± SD of cell area (n = 15 cells) in response to hypo-osmotic stimuli (grey rectangle indicate the treatment, 120 s of switch from TAP (black line, Hypo) or from TAP + 100 mM sorbitol (yellow line, Hypo +) to deionized water + 0.34 mM $CaCl_2$). Significantly different values from the control (untreated cells, CTR) are marked with different letters indicating the treatment (a, Hypo; b, Hypo +) as determined by One-way ANOVA ($P < 0.05$). b. Normalized cell area variations (referred to a.) after the perfusion was restored to the starting medium following hypo-osmotic shock (200 s - 300 s values). c. Normalized FRET Ratio trace and cell area variation of a single representative cell upon hypo-osmotic stimulus (Hypo +, grey rectangle).

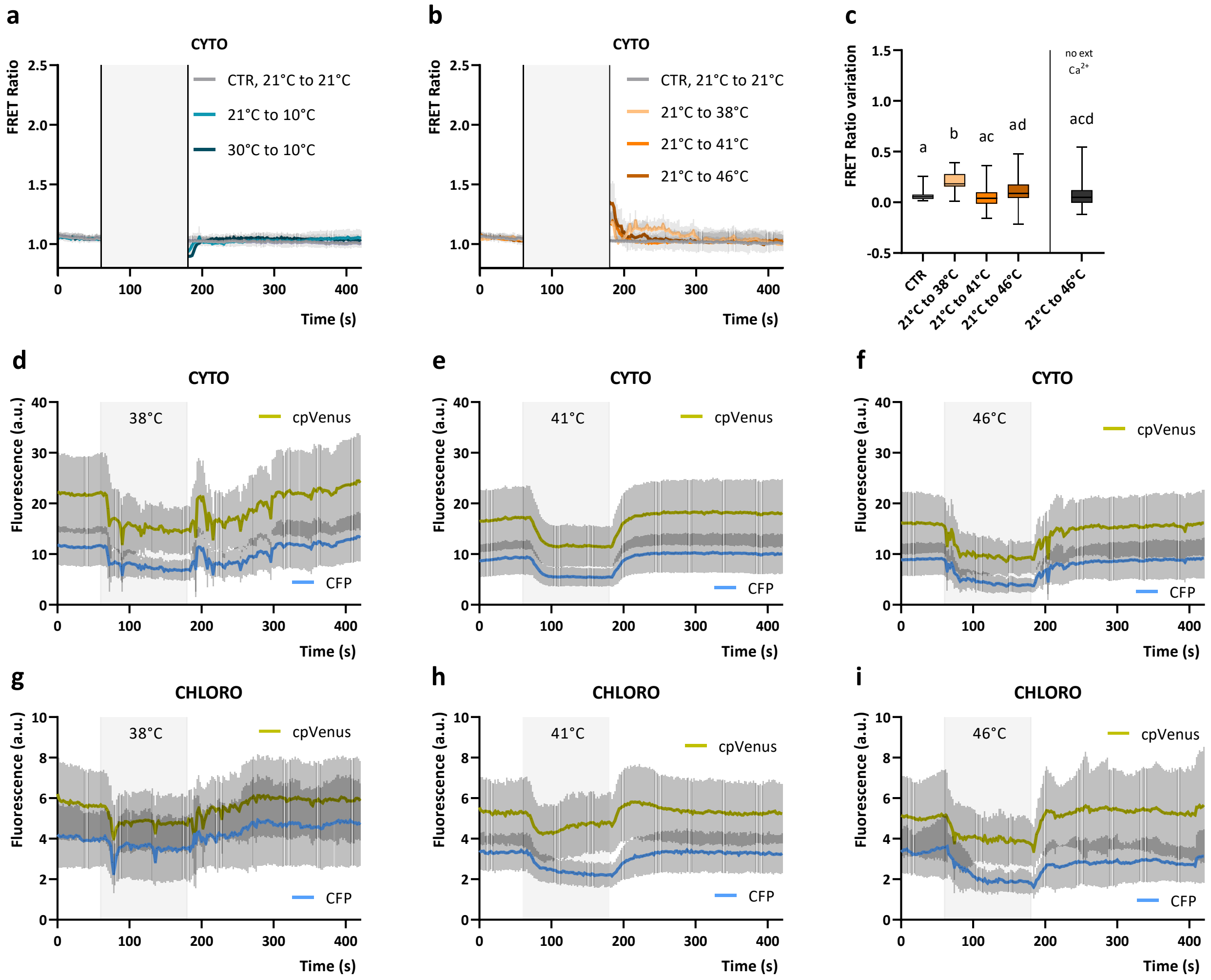


**Figure S4. Compartment-specific $Ca^{2+}$ dynamics in *C. reinhardtii* cells treated with elevated temperatures.** a, b. Averaged and normalized FRET Ratio ± SD of cytosolic YC3.6 in C. *reinhardtii* cells in response to cold shock (a, n > 39 cells) and heat stress (b, n > 32 cells) (grey rectangle indicate the treatment, 120 s of switch of perfusion at different temperature, 21°C-21°C (CTR), 21°C-10°C, 30°C-10°C, 21°C-38°C, 21°C-41°C, 21°C-46°C). c, Maximum FRET Ratio variations after stimulus triggered by heat stress in the cytosol (referred to b). One-way ANOVA: P value < 0.05. d, e, f. Reported in the figures are single wavelength emission of cpVenus (yellow trace) and CFP (light blue trace) of the YC3.6 probe used for the ratio calculations in **Errore. L'origine riferimento non è stata trovata.**b d, e, f; CYTO) and **Errore. L'origine riferimento non è stata trovata.**4b (g, h i; CHLORO) in response different heat stress treatments. The grey rectangle in each graph indicates the treatment at the indicated final temperature. The decrease of both CFP and cpVenus observed upon heat stress treatment is likely due to a temperature-dependent transient change of YC3.6 fluorescence emission properties.

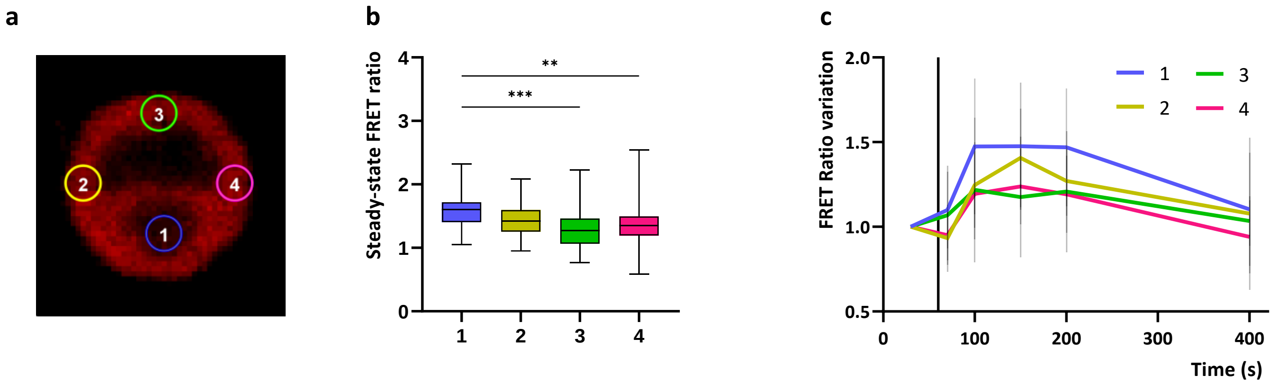


**Figure S5. FRET ratios at the level of the different subchloroplast regions of cells grown in photoautotrophic conditions (TP, minimal medium).** a. Chlorophyll fluorescence image reporting the drawn regions of interest (ROIs) considered at the level of the different subchloroplast regions. b. Basal steady-state FRET ratios at the level of the different subchloroplast regions of photoautotrophic grown cells. (numbers refer to ROIs of a.; n = 47 cells). One-way ANOVA: ***, $P < 0.001$; **, $P < 0.01$. c. Averaged and normalized subchloroplast ROIs FRET ratio variations ± SD (colors refer to a., n = 10 cells examined where the pyrenoid region could be clearly distinguished) in response to 40 mM sodium bicarbonate (black line indicates the treatment onset, switch from growth medium to growth medium + 40 mM $NaHCO_3$). One-way ANOVA ($P > 0.05$).

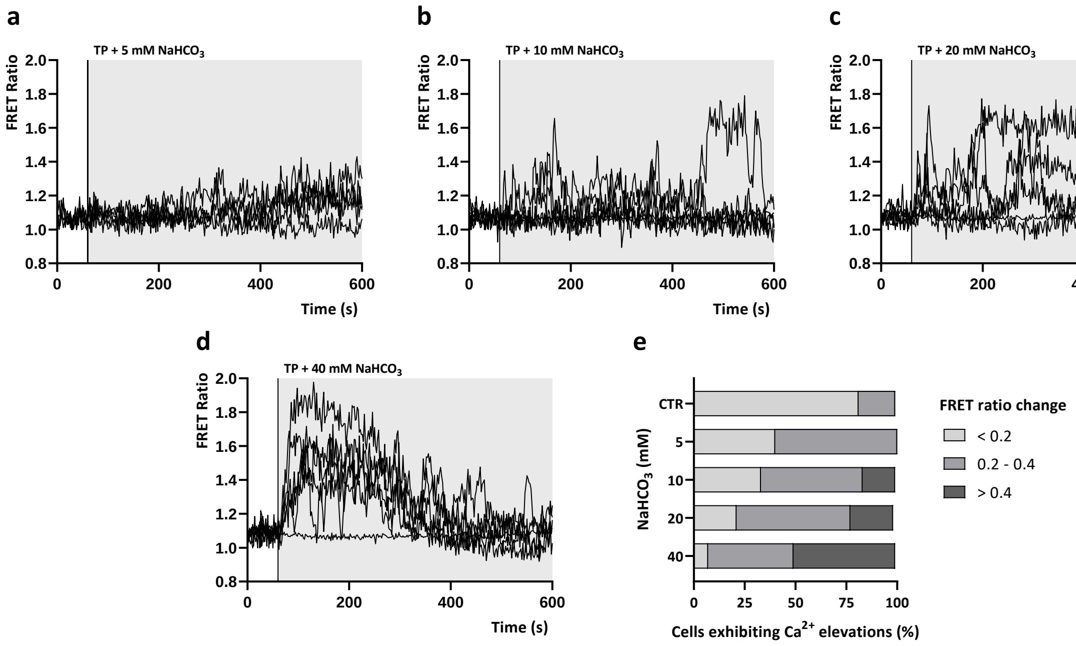


**Figure S6. Stimulus-specific chloroplast $Ca^{2+}$ transients in response to $NaHCO_3$ treatment**. a-d. Normalized chloroplast FRET ratio traces of five representative cells in response to different $NaHCO_3$ concentrations in TP medium, pH 7 (black vertical line indicates the onset of the stimulus, 60 s). e. The percentage of cells exhibiting different FRET ratio elevations in response to specific $NaHCO_3$ treatments (n > 10 cells).

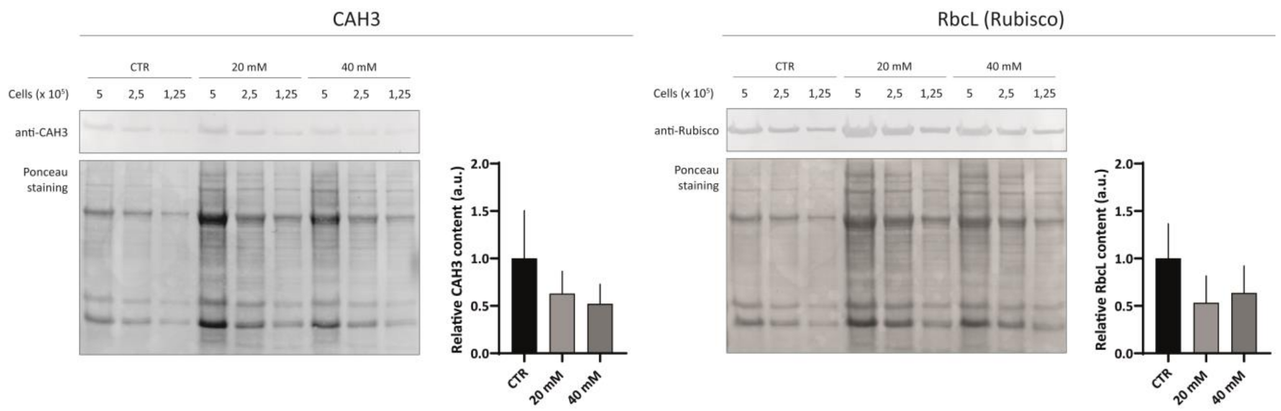


**Figure S7. CAH3 and RbcL protein expression levels in autotrophically grown *C. reinhardtii* cells.** Reported are images of the western blot analysis (upper panel, using specific antibodies anti-CAH3 and anti-RbcL) and Ponceau staining (lower panel) used for the immunotitration of CAH3 and RbcL protein accumulation (graphs in the right side). Lanes were loaded with the total cell amount indicated of cultures daily supplemented with different sodium bicarbonate concentrations (CTR, no supplement; 20 mM sodium bicarbonate; 40 mM sodium bicarbonate). Data were corrected for total protein amount, considering Ponceau stained lanes as reference, and normalized to the control (CTR, 1). Data in the graphs are expressed as Mean ± SD (n = 3). One-way ANOVA: P value > 0.05.

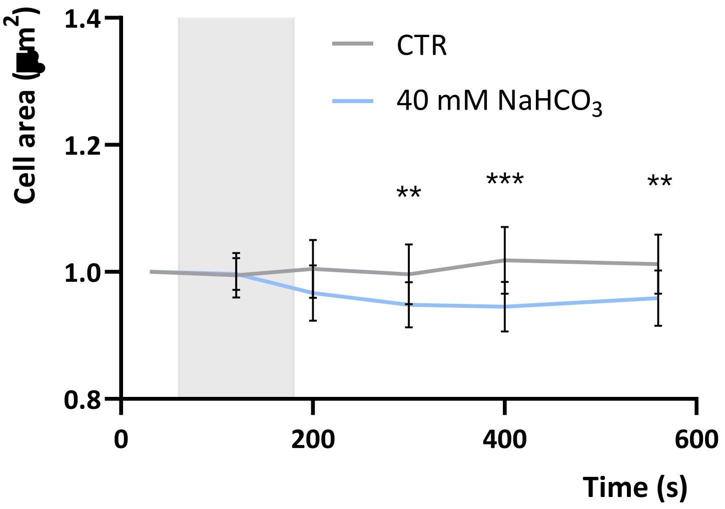


**Figure S8.** ***C. reinhardtii*** **cell area variation in response to 40 mM NaHCO$_3$.** Averaged and normalized cell area measurements ± SD of cell area (n = 15 cells) in response to 40 mM NaHCO$_3$ (grey rectangle indicate the treatment, 120 s of switch from growth medium to growth medium + 40 mM NaHCO$_3$,). Significantly different values from the control (CTR, untreated cells, from figure S2) are marked with asterisks: multiple t-test analysis (**, $P < 0.01$; ***, $P < 0.001$).
